# The novel Rab5 effector FERRY links early endosomes with the translation machinery

**DOI:** 10.1101/2021.06.20.449167

**Authors:** J. S. Schuhmacher, S. tom Dieck, S. Christoforidis, C. Landerer, J. Davila Gallesio, L. Hersemann, S. Seifert, R. Schäfer, A. Giner, A. Toth-Petroczy, Y. Kalaidzidis, K. E. Bohnsack, M. T. Bohnsack, E. M. Schuman, M. Zerial

## Abstract

Localized translation is vital to polarized cells and requires precise and robust distribution of different mRNAs and ribosomes across the cell. However, the underlying molecular mechanisms are poorly understood and important players are lacking. Here we show that the novel Rab5 effector <u>F</u>ive-subunit <u>E</u>ndosomal <u>R</u>ab5 and <u>R</u>NA/ribosome intermediar<u>Y</u>, FERRY complex recruits mRNAs and ribosomes to early endosomes, through direct mRNA interaction. FERRY displays preferential binding to certain groups of transcripts, including mRNAs encoding mitochondrial proteins. Deletion of FERRY subunits reduces the endosomal localization of transcripts in cells and has a significant impact on mRNA and protein levels. Clinical studies show that genetic disruption of FERRY causes severe brain damage. We found that, in neurons, FERRY co-localizes with mRNA on early endosomes and mRNA loaded FERRY-positive endosomes are in close proximity of mitochondria. FERRY thus transforms endosomes into mRNA carriers and plays a key role in regulating mRNA distribution and transport.

## Introduction

Subcellular mRNA localization and protein translation is vital for fundamental biological processes, such as embryonic development, cellular homeostasis, neuronal plasticity and adaptive response to environmental cues (Cioni et al., 2018; Das et al., 2021; Glock et al., 2017; Martin and Ephrussi, 2009; Turner-Bridger et al., 2020). While the asymmetric localization of specific mRNAs during oogenesis (Becalska and Gavis, 2009; Riechmann and Ephrussi, 2001) represents a morphologically simple example, a completely different scenario unfolds in the brain, where neurons span long distances with their axonal and dendritic processes. Not only are these compartments highly specialized in their function, but they also respond to external cues on a millisecond timescale at the distal end of their network, far away from the cell body. Neurons handle these challenges by producing several proteins at their site of action through local translation, which is involved in axon outgrowth, branching synaptogenesis, regeneration and neuronal plasticity (Cioni et al., 2018; Jung et al., 2014; Kim and Jung, 2020; Rangaraju et al., 2017). Local translation implies the availability of the mRNAs at the sites of respective protein function, and hence the precise subcellular localization of a plethora of mRNAs (Glock et al., 2017; Turner-Bridger et al., 2020).

The correct transport and subcellular localization of mRNAs requires a sophisticated molecular regulation tailored to the specific roles of the mRNAs and their encoded products. Transcriptomic studies have identified thousands of different mRNAs in neuronal sub-compartments, such as axons, dendrites or the neuropil (Andreassi et al., 2010; Briese et al., 2016; Cajigas et al., 2012). Furthermore, these transcripts are distributed heterogeneously with mRNAs showing distinct localization patterns, for example, being restricted to axons or dendrites or even smaller sub-compartments. These findings reflect the existence of a complex mRNA distribution plan where thousands of mRNAs have to find their correct location.

Such a complex task and long distances, especially in neurons, are incompatible with a passive diffusion-based mechanism and require active mRNA transport along the cytoskeleton. A direct connection between RNA-binding proteins (RBPs) and motors proteins has been observed in various forms, for example the targeting of mRNAs by RBPs that recognize *cis*- regulatory elements on the respective mRNA, including the so called ‘zipcodes’ (reviewed in: (Buxbaum et al., 2015; Das et al., 2021)). Recently, different compartments of the endolysosomal system have been associated with the spatial organization of components of the translation machinery, including mRNAs, mRNP granules and ribosomes in various organisms (Cioni et al., 2019; Higuchi et al., 2014; Liao et al., 2019). The endolysosomal system acts as a central logistic system of eukaryotic cells, comprising multiple membrane-enclosed organelles, such as early endosomes (EE), late endosomes and lysosomes, which traffic and sort a large variety of cargos.

It is ideally suited to regulate mRNA transport and localization, especially in morphologically complex and spatially segregated compartments, like the hyphae of fungi or the processes of neurons. In the fungus *U. maydis*, a special adaptor system enables the long-distance transport of mRNAs and polysomes on EEs (Higuchi et al., 2014). In higher eukaryotes, lysosomes serve as an Annexin A11-mediated mRNP granule transport vehicle, while late endosomes act as translation platforms for mitochondrial proteins in neurons (Cioni et al., 2019; Liao et al., 2019).

A recent study reported the co-localization of mRNAs to EEs, suggesting that they may also be part of an mRNA distribution machinery (Popovic et al., 2020). The EE is an early sorting station for cargos coming from the plasma membrane, which are routed towards recycling or degradation. EEs appear more suitable to support directional mRNA transport than late endosomes, due to their bidirectional motility in neurons (Goto-Silva et al., 2019) whereas late endosomes (multi-vesicular bodies) primarily migrate retrograde (Parton et al., 1992). The identity of endosomes is determined by an intricate interplay between proteins and specific lipids that are intimately linked to Rab GTPases (Pfeffer, 2013; Wandinger-Ness and Zerial, 2014). Different Rab GTPases characterize different endocytic organelles, such as Rab4 and Rab11 recycling endosomes and Rab7 late endosomes (reviewed in: (Wandinger-Ness and Zerial, 2014)). Rab5 is the hallmark GTPase of the EE and a membrane organizer. Upon activation from the GDP- to the GTP-bound form on the EE, Rab5 recruits a plethora of Rab5 effectors, such as the molecular tether EEA1 (Christoforidis et al., 1999) or Rabankyrin-5 (Schnatwinkel et al., 2004), thereby orchestrating different functions of the organelle (Cezanne et al., 2020; Franke et al., 2019; Lauer et al., 2019; Lippe et al., 2001; Murray et al., 2016). To date, the molecular mechanism describing the connection between EEs and mRNAs or the translation machinery remains mysterious. No known mRNA-associated protein appears to localize on EEs nor do any endosomal proteins exhibit classical RNA-binding motifs. Considering the large number of precisely localized mRNAs, a versatile molecular machine able to discriminate between different mRNAs and transport specific mRNA subgroups would be efficient and only use a limited number of carriers.

Closing this gap, we report the discovery of a novel five-subunit Rab5 effector complex, which we named <u>F</u>ive-subunit <u>E</u>ndosomal <u>R</u>ab5 and <u>R</u>NA/ribosome intermediar<u>Y,</u> FERRY complex. Through direct interaction with Rab5 and mRNAs, it connects the EE with the translation machinery and is important for mRNA distribution and transport.

## Results

### Identification of a novel Rab5 effector complex

In previous studies, we isolated Rab5 effectors using a Rab5 affinity chromatography (Christoforidis et al., 1999). Upon further purification of this elaborate set of proteins, we observed five proteins co-fractionating in size exclusion chromatography (SEC) (Figure S1A, left panel). Further purification using ionic charges resulted in co-elution of the same set of five proteins (Figure S1A, right panel). The co-fractionation during both chromatography steps indicated that these proteins form a complex, raising a great interest regarding its identity and function. Mass spectrometry revealed the five proteins as Tbck (101 kDa), Ppp1r21 (88 kDa), C12orf4 (64 kDa), Cryzl1 (39 kDa) and Gatd1 (23 kDa) (Figure 1A). For clarity, we will refer to the novel complex as the <u>F</u>ive-subunit <u>E</u>ndosomal <u>R</u>ab5 and <u>R</u>NA/ribosome intermediar<u>Y</u> (FERRY) complex, with the individual subunits being designated Fy-1 – Fy-5 (Figure 1A).

**Figure 1:**
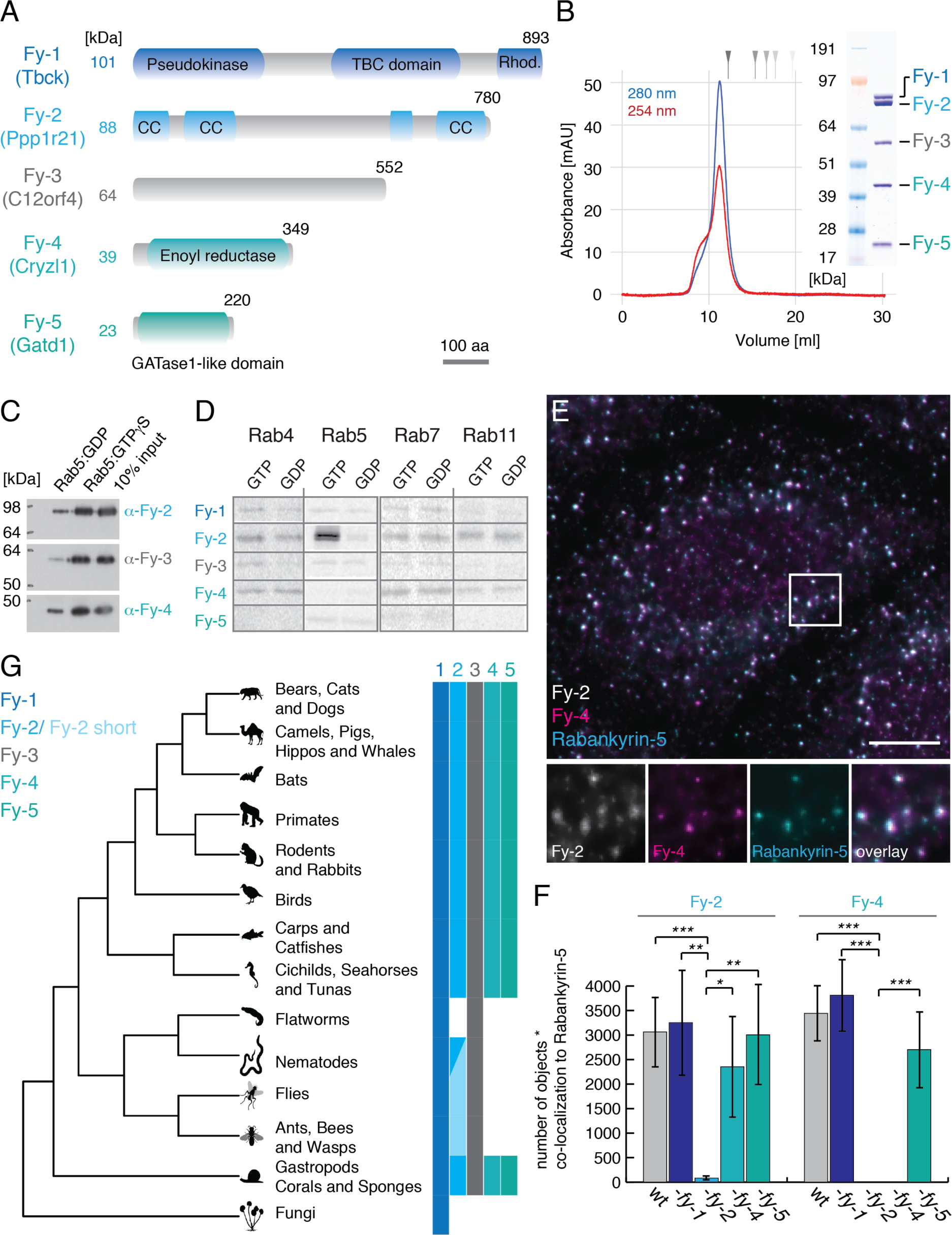
**A)** Domain architecture of the components of the FERRY complex drawn to scale (scale bar in left lower corner: 100 amino acids (aa); TBC: Tre-2/Bub2/Cdc16, Rhod.: Rhodanese domain, CC: coiled-coil). **B)** SEC profile of the FERRY complex (blue: 280 nm, red: 254 nm) with a Coomassie-stained SDS PAGE. Molecular weight standard (670, 158, 44, 17, 1.35 kDa). **C)** Western blot analysis of an *in vitro* pulldown assay of the FERRY complex incubated with glutathione beads with GST-Rab5 loaded with GDP or GTPγS using antibodies against Fy-2, Fy-3 and Fy-4. **D)** Fluorographic analysis of GST binding assays using different Rab GTPases against *in vitro* translated ^35^S methionine-containing FERRY components. **E)** Immunostaining of HeLa cells against Rabankyrin-5, Fy-2 and Fy-4 (Scale bar: 10 μm). The boxed region is shown below in more detail. **F)** Quantification of the fluorescent signal of Fy-2 and Fy-4 in images as in E. **G)** Phylogenetic analysis of the subunits of the FERRY complex (complete list: Table S1).

In a first step, we successfully reconstituted the FERRY complex *in vitro*. Figure 1B shows that the five proteins form a stable complex, as they all elute as a single peak from SEC. To estimate the stoichiometry of the components in the complex, we compared the intensity of the corresponding signals of a Coomassie-stained SDS PAGE, which suggested a ratio of 1:2:1:2:4 for Fy-1:Fy-2:Fy-3:Fy-4:Fy-5, respectively. Using mass photometry, we obtained a molecular weight of 525 ± 41 kDa for the FERRY complex which fits very well with the estimated ratios and a calculated molecular weight of 521 kDa (Figure S1B). This was further corroborated by a cryoEM structure which showed a ratio of 2:2:4 for Fy-2, Fy-4 and Fy-5 (Quentin et al., 2021). With the FERRY complex in hand, we tested whether it fulfills the typical criterion of Rab5 effectors and binds predominantly to the activated, GTP-loaded Rab5, by performing a Glutathione-S-transferase (GST) pulldown assay. Sodium dodecylsulfate polyacrylamide gel electrophoresis (SDS PAGE) and Western blot analysis of different FERRY subunits (Fy-2, Fy-3, Fy-4) revealed a much stronger signal for Rab5:GTPγS than Rab5:GDP, indicating that the FERRY complex interacts preferentially with activated Rab5 (Figure 1C, Figure S1D).

We next tested the specificity of the FERRY complex subunits for different endosomal Rab GTPases, by performing binding assays of *in vitro* translated, ^35^S methionine labelled Fy-1 to Fy-5 against Rab5, Rab4, Rab7 and Rab11 (Figure 1D). In this experimental set up, the binding of each component of the complex was tested individually, in the absence of the other subunits, thereby allowing identification of the subunit(s) of the complex that mediate binding between the FERRY complex and Rab5. Out of the five subunits, only Fy-2 bound to Rab5:GTP, but not Rab5:GDP (Figure 1D). In addition, no interaction was observed between the FERRY complex and the other Rab GTPases, neither in the GDP- nor GTP-bound form. These results indicate that Fy-2 mediates the interaction between the FERRY complex and Rab5:GTP, but none of the endosomal Rab GTPases tested. This was also confirmed by hydrogen deuterium exchange mass spectrometry (HDX-MS), which identified the Rab5 binding site of the FERRY complex near the C-terminus of Fy-2 (Quentin et al., 2021). These results indicate that the FERRY complex is indeed a Rab5 effector.

### The FERRY complex localizes to EEs

The aforementioned specificity of the FERRY complex for Rab5 suggests that it may localize to EEs. The localization of endogenous FERRY complex requires antibodies suitable for immunofluorescence. We were able to raise antibodies against the subunits Fy-2 and Fy-4 that are suitable for immunofluorescence (Figure S1C, see also Methods: Antibody validation). The overall appearance of the fluorescence signal of Fy-2 and Fy-4 revealed a punctate localization pattern in HeLa cells that resembles the distribution of EEs (Figure 1E). As expected, Fy-2 co- localize very well with Fy-4 (0.85) but also with the early endosomal markers Rabankyrin-5 (0.87) and EEA1 (0.76) (Figure S1E), suggesting that the FERRY complex localizes to EEs. To determine whether the FERRY complex is indeed a stable protein complex in cells, we generated HeLa knock-out (KO) cell lines of the FERRY subunits Fy-1, Fy-2, Fy-4 and Fy-5 using CRISPR/Cas9 technology. Loss of the respective protein was confirmed by Western blot analysis (Figure S1F), which also showed that the levels of Fy-3 were reduced upon *fy-2* KO (80%) and *fy-1* KO (20%) (Figure S1F). Subsequently, we assessed the localization of Fy-2 and Fy-4 under these conditions by counting the number of fluorescent structures co-localizing with the EE marker Rabenkyrin-5. The localization of Fy-2 was not significantly changed, except in the *fy-2* KO cell lines. However, in case of Fy-4 we observed a complete loss of EE co-localization in the *fy-2* and *fy-4* KO cell lines (Figure 1F). This is in agreement with biochemical and structural data that identified Fy-2 as mediator of the FERRY Rab5 interaction and, thus to the EE.

The five FERRY subunits exhibit a substantial variability in size, domain composition and structural features. Indeed, the FERRY complex does not resemble any known endosomal complex (*e.g.* CORVET/HOPS, or the ESCRT) (Figure 1A). Searching for traces of the FERRY complex in the course of evolution, we performed a phylogenetic analysis of the subunits of the FERRY complex. While Fy-1 is the most ancestral subunit with homologues in some fungi, we also found an assembly of Fy-1, Fy-3 and a short version of Fy-2 in insects and some nematodes. With the evolution of the Chordata, we observed a transition from a 3- component assembly to the five-subunit complex, via the co-occurrence of two novel proteins, Fy-4 and Fy-5 and the extension of Fy-2 with the Fy-4 and Fy-5 binding sites (Figure 1G, Table S1). This co-evolution further supports the formation of a complex by the FERRY subunits.

### The FERRY complex associates with the translation machinery

Even though the FERRY complex has not previously been identified, it may play an important role in brain function. Clinical studies on patients with mutations in the *fy-1* or *fy-2* genes, showed that loss of either of these proteins severely impairs brain development and function, causing symptoms such as a mental retardation, intellectual disability, hypotonia, epilepsy, and dysmorphic facial features resulting in premature death of the patients (Bhoj et al., 2016; Chong et al., 2016; Guerreiro et al., 2016; Hancarova et al., 2019; Loddo et al., 2020; Ortiz-Gonzalez et al., 2018; Philips et al., 2017; Suleiman et al., 2018; Zapata-Aldana et al., 2019). Different studies report the accumulation of lipofuscin in the human brain and further indicate disturbances in the endocytic system (Beck-Wodl et al., 2018; Rehman et al., 2019). These results suggest that the FERRY complex carries out an endocytic function essential for brain development and neuronal function.

To gain insights into the cellular role of the FERRY complex, we examined its interaction network using a GST pulldown approach (Figure 2A). In a first step, we purified a GST fusion variant of the FERRY complex (GST-FERRY, Figure S2A). Subsequently, GST-FERRY was incubated with fresh HEK 293 cell lysate (see Methods: HEK 293 lysate preparation), stringently washed and eluted from the resin. Mass spectrometry of the elution fractions revealed 34 potential interaction partners of the FERRY complex (Figure 2B, Table S2). Almost three-quarters of the candidates (73.5%) represent ribosomal proteins of both the large and the small subunit (Figure 2C), suggesting that complete ribosomes and maybe the translation machinery may be associated with the FERRY complex. Due to their abundance, ribosomal proteins are frequent contaminants of such assays. To further test the ribosome association of FERRY, we generated stably transfected HEK293 cell lines in which expression of Flag-His-Fy-2 or Fy-2-His-Flag can be induced. Subsequently, cell lysates were fractionated using a sucrose gradient separating the small and the large ribosomal subunits, monosomes and polysomes from smaller complexes and free proteins and RNAs (Figure S2B). While the majority of Fy-2 was found to be non-ribosome-associated, a fraction of Fy-2 was also observed co-migrating with the different subunits, monosomes and a minor fraction was also detected with polysomes, supporting the hypothesis that the FERRY complex is able to associate with ribosomes in cells (Figure 2D).

**Figure 2:**
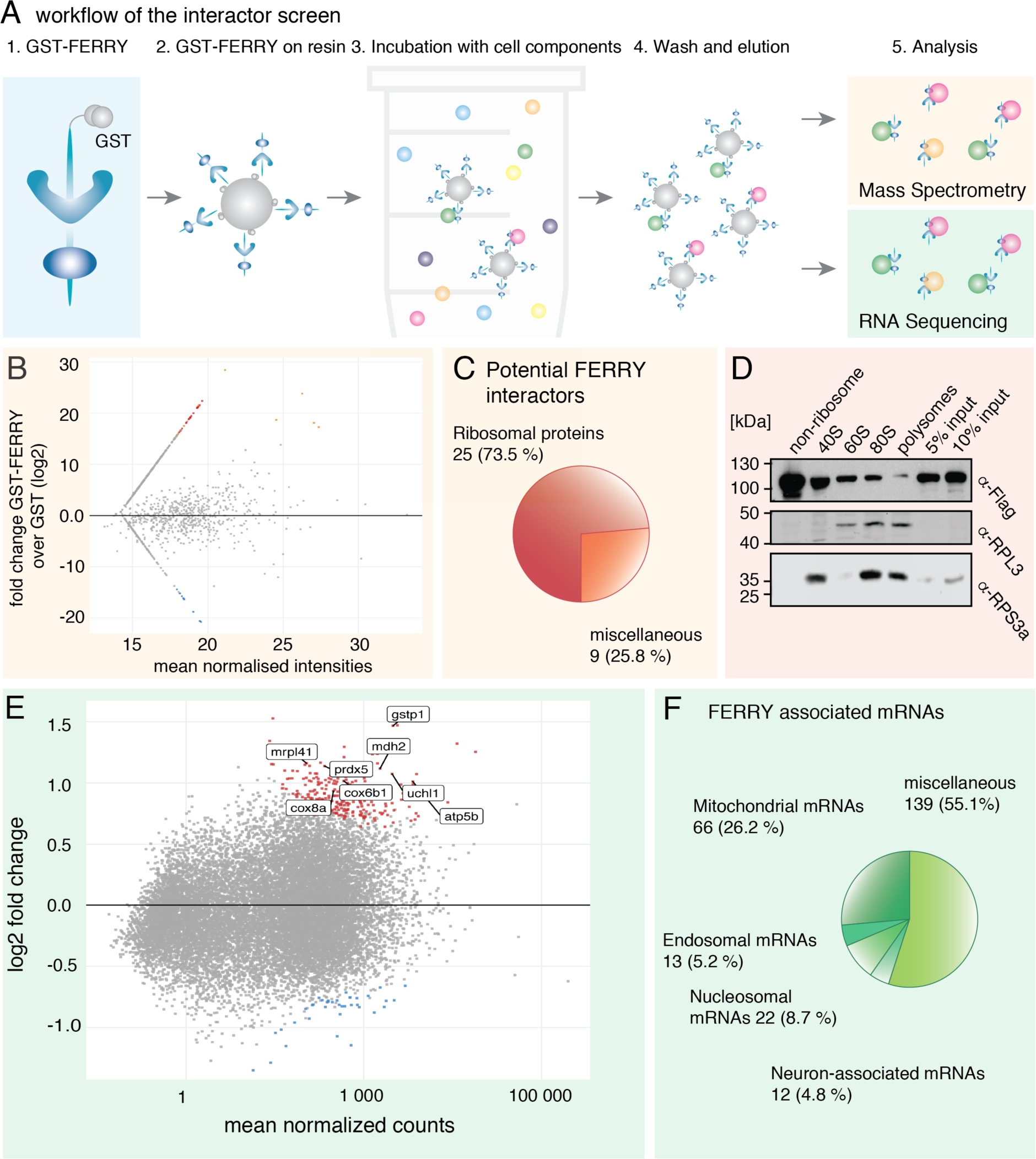
**A)** Scheme of the *in vitro* GST-FERRY interactor screen. **B)** MA blot of results of the GST-FERRY interactor screen. Candidates enriched in GST-FERRY and GST are indicated in red and blue, respectively. **C)** Pie chart of potential FERRY interactors **D)** Western blot analysis of sucrose density gradient fractions containing ribosomal (40S, 60S, 80S and polysomes) and non-ribosomal complexes **E)** MA blot of the RNA sequencing of potential FERRY-associated mRNAs. mRNA candidates associated with GST-FERRY and GST are highlighted in red and blue, respectively **F)** Pie chart of the FERRY-associated mRNAs.

The association of the FERRY complex with ribosomes prompted us to assess the spectrum of FERRY interactors further. We tested whether RNAs accompany the ribosomes and RNA-binding proteins as FERRY interactors. To identify transcripts co-eluting with the FERRY complex, we modified the protocol of the GST-FERRY pulldown assay to obtain RNA instead of proteins, which was subsequently analyzed by sequencing (Figure 2A). Applying a stringent cut-off (adjusted p-value (padj) < 0.01), the experiment revealed 252 mRNAs significantly associated with the FERRY complex (Figure 2E, Table S2). Among these candidates, the largest group of mRNAs (66 transcripts/ 26.2%) constitute nuclear-encoded mitochondrial proteins. Furthermore, we also identified components of the endosomal system and nucleosome components (Figure 2F). A gene set enrichment analysis against a gene set collection (MSigDB C5 collection: ontology gene sets), revealed a strong enrichment for mitochondrial matrix genes (#1714), mitochondrial ribosome (#2354), cellular respiration (#480) and TCA cycle (#4413) components. In summary, these results suggest that the FERRY complex interacts with specific groups of mRNAs, especially those encoding mitochondrial proteins.

### The FERRY complex interacts directly with mRNA

To test the hypothesis of a direct FERRY interaction with mRNA, we performed electrophoretic mobility shift assays (EMSA) with *in vitro* transcribed mRNAs. Initially, we chose *mrpl41,* a top candidate of the RNA screen and included the 5’ untranslated region (UTR), the open reading frame (orf), the 3’-UTR and a short stretch of 50 adenines, yielding a 660-nucleotide, artificially poly-adenylated mRNA. With increasing amounts of *mrpl41* mRNA an additional signal at a higher molecular weight appeared in the EMSA, indicating a binding of the FERRY complex to the *mrpl41* mRNA (Figure 3A).

**Figure 3:**
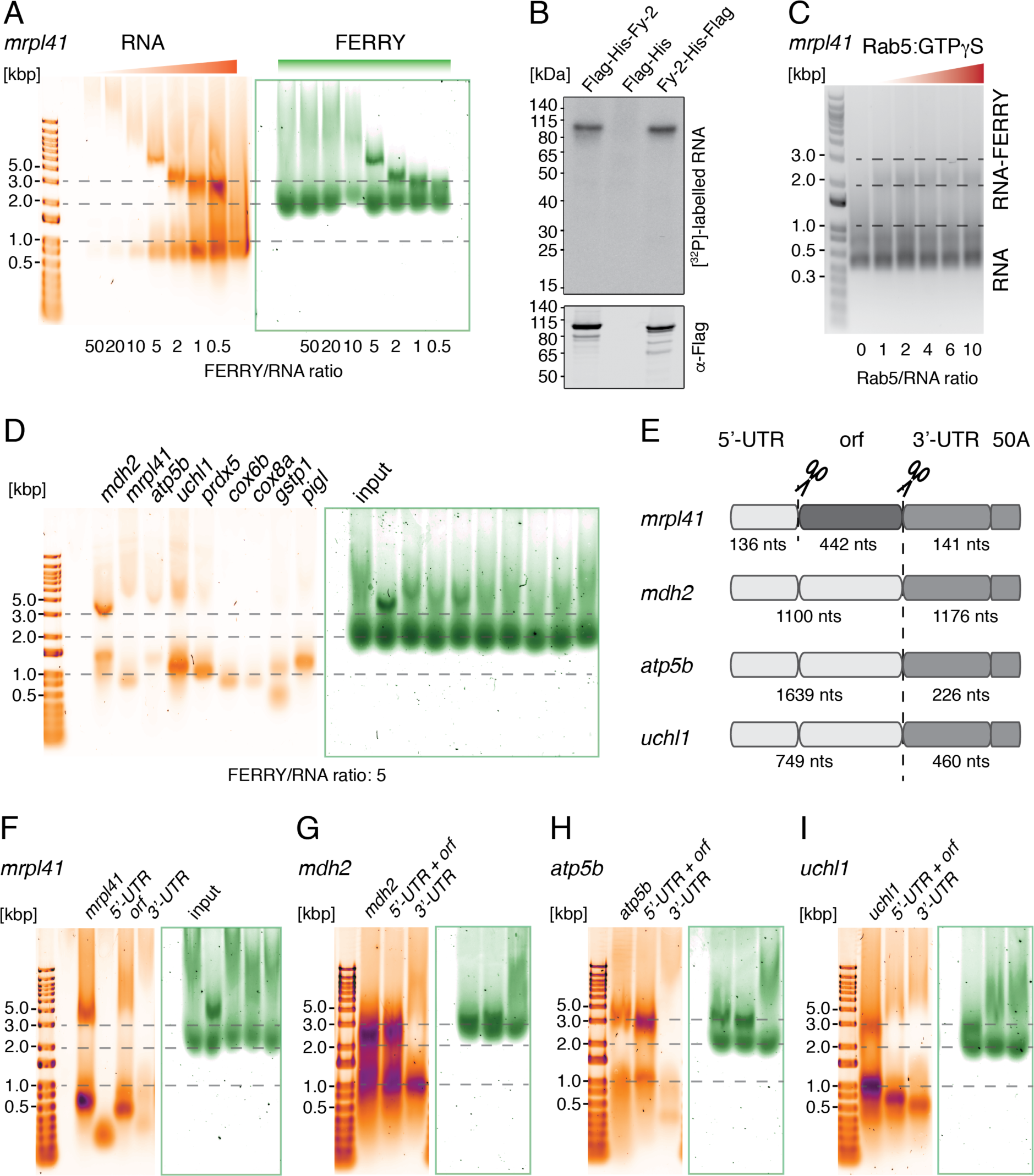
**A)** Electrophoretic mobility shift assay (EMSA) with increasing ratios of FERRY to RNA. (RNA: orange, SYBR Gold; proteins: green, Sypro Red) **B)** Detection of radiolabelled RNAs by autoradiography and Flag-tagged proteins by Western blotting of a tandem affinity purification after UV crosslinking. **C)** EMSA in the presence of Rab5:GTPγS with a fixed ratio of FERRY to RNA of 3. (RNA: grey, ethidium bromide) **D)** EMSA with different mRNAs at a fixed FERRY/RNA ratio of 5. **E)** Scheme of RNA sub-constructs. **F) – I)** EMSAs comparing four different RNAs with their respective subdivision construct shown in E.

Next, we aimed at investigating whether the FERRY complex also directly interacts with RNA in cells. Therefore, we used UV-mediated protein-RNA cross-linking, which is a zero-distance cross-linking method that covalently attaches proteins to bound RNAs. We utilized the two HEK293 cell lines expressing Flag-His-Fy-2 and Fy-2-His-Flag, as the main RNA interface of the FERRY complex is located on Fy-2 (Quentin et al., 2021). After cross-linking, we isolated the Fy-2 tagged proteins by tandem affinity purifications under native (anti-Flag) and strongly denaturing (Ni^2+^ affinity) conditions and confirmed their correct size by Western blot (Figure 3B). Furthermore, the isolated material was examined regarding the presence of cross-linked RNA. RNA visualization using ^32^P labelling after partial RNase digestion revealed a signal at the correct molecular weight for both Fy-2 variants, while the control lane with only the His-Flag sample was empty (Figure 3B). This experiment confirms a direct FERRY-RNA interaction in cells, which is mediated by Fy-2.

We further characterized the interaction using EMSA assays by testing whether the binding of the model mRNA *mrpl41* to the FERRY complex is Rab5-dependent. We performed EMSAs with a fixed FERRY/*mrpl41* mRNA ratio and added increasing amounts of Rab5:GTPγS to the assay. This did not have a visible effect on the FERRY-mRNA interaction, suggesting that Rab5 does not play a role in this process (Figure 3C).

The enrichment of specific subsets of mRNAs in the RNA screen points towards the ability of the FERRY complex to discriminate between different mRNAs. To examine the specificity of mRNA binding, we chose eight mRNAs from the 237 found in the screen, that encode proteins fulfilling different mitochondrial functions, such as components of the respiratory chain (*cox6b* and *cox8a*), the ATP Synthase (*atp5f1b*), the mitochondrial stress response (*gstp1* and *prdx5*), the mitochondrial ribosome (*mrpl41*), the TCA cycle (*mdh2*) and the mitochondrial ubiquitination machinery (*uchl1*), and tested their interaction with the FERRY complex using EMSAs. We additionally included the mRNA of *pigl* as a negative control, as this mRNA neither appeared enriched in the GST-FERRY pulldown assay, nor was significantly changed in the transcriptome analysis of FERRY KO cell lines (see below). While *mrpl41*, *mdh2* and *atp5f1b* exhibited a clear interaction with the FERRY complex, the interaction with the other five candidates was much weaker (Figure 3D, Figure S3A). These results suggest that the FERRY complex binds transcripts with different efficacy *in vitro*.

To further establish to which classes of RNAs the FERRY complex preferable binds, we tested its ability to interact with small RNAs (< 200 nts) in general and different tRNAs (tRNA^Arg(ACG)^, tRNA^Cys(GCA)^ and tRNA^Phe(GAA)^)) more specifically using EMSA assays. Even at equimolar FERRY-to-RNA ratios, we were not able to detect any interaction, indicating a certain preference of FERRY for mRNAs (Figure S3B-D).

Although we selected relatively short mRNA candidates for our *in vitro* assays, these RNAs still have considerable lengths, ranging from 600 to 2200 nucleotides. This raises the question regarding the region on the RNA to which the FERRY complex binds. We therefore chose four mRNAs (*mrpl41*, *mdh2*, *atp5b* and *uchl1*) showing clear binding to the FERRY complex and divided them into different parts (Figure 3E). The mRNA of Mrpl41 was split into three parts, the 5’-UTR, the orf and the 3’UTR with an addition of 50 adenine nucleotides. While the two UTR fragments did not show any interaction with the FERRY complex, the orf fragment was still able to bind FERRY. However, the interaction was weaker than observed with full length *mrpl41* mRNA (Figure 3F, Figure S3E). The other three candidates were each split into two parts, the 5’-UTR + orf and the 3’-UTR with 50 adenines. The *mdh2*-FERRY and *atp5b*- FERRY interactions were clearly mediated by the 5’-UTR + orf fragments, while the 3’-UTR + 50A fragments did not bind the FERRY complex (Figures 3GH, Figures S3FG). Interestingly, for *uchl1* mRNA, both parts still interacted with the FERRY complex, albeit showing reduced binding (Figure 3I, Figure S3H). Altogether, these results imply that the FERRY complex does not bind to a single, short motif on the mRNAs. This conclusion is supported by the structural analysis of FERRY bound to mRNA, which showed a large and complex interface involving different subunits of the FERRY complex (Quentin et al., 2021). Such an extended binding interface clearly distinguishes the FERRY complex from other RBPs connected to mRNA transport, such as ZBP1, FMRP, Staufen2 or the proteins of the elavl family, that bind to distinct, mostly AU-rich motifs, in the 3’-UTR (Schieweck et al., 2020).

### The FERRY complex impacts mRNA localization in HeLa cells

To investigate the cellular role of the FERRY-mRNA interactions, we designed an experiment to compare the localization of EEs (marked by EEA1) and different mRNAs, including a probe against *polyA* that reflects the general mRNA distribution (Figure 4A). In *wildtype* (*wt*) cells, we quantified a 13.3% co-localization between *polyA*, mRNA and EEA1-positive EEs (Figure 4B, C).

**Figure 4:**
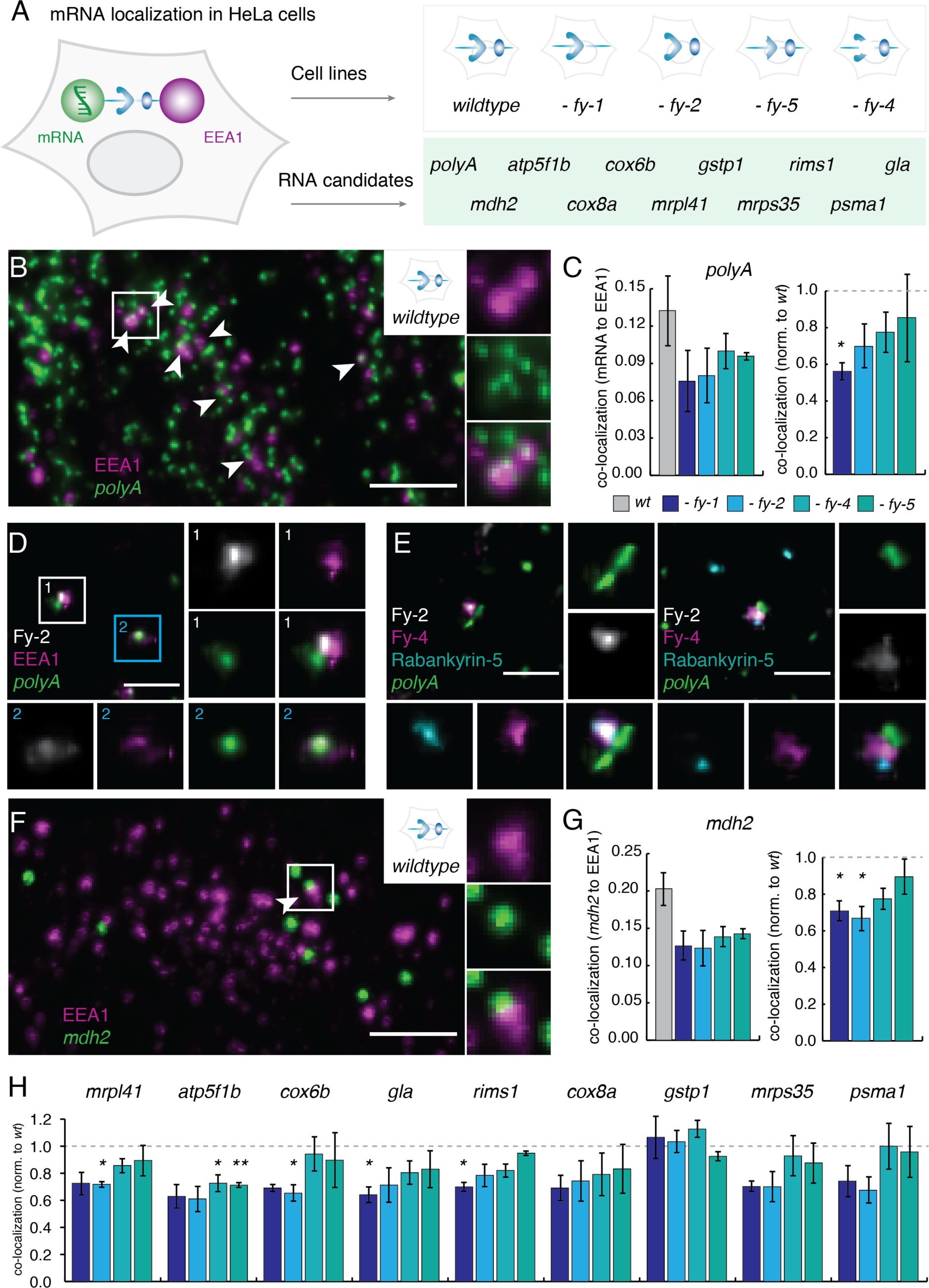
**A)** Scheme of the localization experiment, showing different markers (mRNA: smFISH, EEA1: antibody), mRNAs (in the green box) and cell lines (grey box). **B)** Visualization of EEA1 and *polyA* in *wt* HeLa cells (Scale bar: 5 μm). Co-localization events are indicated with white arrow heads and the boxed region is highlighted on the right. **C)** Quantification of co-localization of *polyA* and EEA1 in HeLa *wt* and different KO cell lines. **D)** and **E)** Super-resolution (Musical) imaging with indicated markers (Scale bar: 1 μm). **F)** Visualization of EEA1 and *mdh2* mRNA in *wt* HeLa cells (Scale bar: 5 μm). The Co-localization event is indicated with an arrow head and the boxed region is highlighted on the right. **G)** Quantification of co-localization of the *mdh2* mRNA probe and EEA1 in different HeLa cell lines. **H)** Quantification of co-localization of the different mRNAs and EEA1 in FERRY KO cell lines (Co-localization normalized to *wt*).

To ensure that the FERRY-mRNA colocalization to EE is specific, we visualized the FERRY complex, the EE and the mRNA concomitantly by multicolor super resolution microscopy. With multiple signal classification (MUSICAL) we were able to acquire up to four different components and reached a resolution of 60 for mRNA and100 nm for endosomal markers (see also Methods). Firstly, we combined Fy-2, EEA1 (EE marker) and mRNA (*polyA*), and observed co-localization as well as partial co-localization with the fluorescence signals in very close proximity (< 200 nm) (Figure 4D). The enhanced resolution allows resolving the fluorescent signals and we could detect instances where Fy-2 appears to bridge EEA1 and the mRNA (Figure 4D, box). Given the sizes of EEA1 in the extended conformation (>200 nm), of Fy-2 and mRNA (Figure S4A), one cannot expect a precise colocalization of signals on the image. The observed distances are within the expected range of a FERRY-mediated attachment of mRNA to the EE (Figure S4A). Secondly, we used both available FERRY markers (Fy-2 and Fy-4) with Rabankyrin-5 (EE marker) and mRNA (*polyA*). Often mRNA, FERRY and EE partially co-localized within a range of 250 nm and we observed events where FERRY and the EE co-localize while the signal for the mRNA is slightly shifted (Figure 4E). Again, we detected events where both FERRY markers are located between EE and mRNA (Figure 4E, box). These data validate the colocalization of FERRY-RNA interaction by confocal microscopy (Figure 4A-C) and corroborate the notion that the FERRY complex connects the EE with mRNA.

Having established the specificity of the co-localization of *polyA* mRNA and FERRY to EE, we examined the localization of a set of mRNAs to EE and the consequence of removal of different FERRY subunits. The candidate set was chosen to include mRNAs that showed binding to the FERRY complex *in vitro* (*i.e. mdh2, mrpl41* and *atp5f1b*), mRNAs that were identified in our RNA association screen but did not interact with FERRY *in vitro* (*cox8a*, *cox6b* and *gstp1*), and mRNAs that were inconspicuous in both experiments (*mrps35*, *rims1*, *psma1* and *gla*). To obtain reliable statistics for mRNAs that are only a small fraction of *polyA* mRNA, we used automated confocal microscopy to obtain a large sample set. We acquired images visualizing EEA1 and mRNA in *wt* and FERRY component KO HeLa cell lines. From 13.3% co-localization in *wt* cells, the EE mRNA co-localization decreased upon KO of each of the four FERRY subunit KOs tested to a varying degree (Figure 4C, left). The KO of *fy-1* had the strongest effect and reduced the frequency of mRNA-EE co-localization by 44%. In the *fy-2*, *fy-4* and *fy-5* KO cell lines, we also observed a reduction of EE mRNA co-localization by 30%, 22% and 15%, respectively (Figure 4C, right). These results indicate that the FERRY complex contributes significantly to the recruitment of mRNA on EEs, in addition to other RBPs (Schieweck et al., 2020).

Next, we analyzed the co-localization of individual mRNAs with EEs and observed a range of 14.4% to 22.7% co-localization in the *wt* cell line (Figure S4B). Again, we detected a decrease in co-localization upon FERRY subunit KO for certain mRNAs. For example, the co- localization of *mdh2* mRNA with EEs decreased from 20% in *wt*, to 12-14% in the KO cell lines, with the most pronounced effect in the *fy-1* and *fy-2* KO cell lines (Figure 4F, G). In general, the loss of one of the large subunits Fy-1 or Fy-2 had a stronger impact on mRNA localization, with several mRNAs (*i.e. mrpl41*, *cox6b*, *gla* and *rims1*) showing a significant decrease in EE localization upon loss of *fy-1* or *fy-2*. Endosomal localization of *atp5f1b* was affected in all four KO cell lines, however, more pronounced upon *fy-4* or *fy-5* KO (Figure 4H). Also, EE localization of *cox8a*, *mrps35* and *psma1* appeared decreased in the *fy-1* and *fy-2* KO cell lines (Figure 4H). In summary, the KOs of various components of the FERRY complex impacts the recruitment of mRNAs to EEs. Furthermore, the loss of the FERRY complex only affects the co-localization of certain mRNAs with EEs, while others are only moderately affected or unaltered. KO of *fy-1* or *fy-2* had a stronger impact on early endosomal mRNA recruitment than the KO of *fy-4* or *fy-5*, which is also reflected by the clinical data that report severe symptoms for mutations in *fy-1* or *fy-2*. These data suggest that the FERRY complex plays a major role in mRNA recruitment to EEs and most probably mRNA transport. The observation that some mRNAs are more affected than others underlines the notion that the FERRY complex exhibits a preference for certain mRNAs.

### The loss of FERRY impacts the cellular transcriptome and proteome

We hypothesized that mis-localization of mRNAs might have an influence on mRNA levels and, possibly, protein expression. To test our hypothesis, we analyzed the transcriptome of the different FERRY component KO cell lines. Data analysis revealed transcriptomic changes in all four KO cell lines (Figure 5A, Figure S5A). Interestingly, we observed a more pronounced up-regulation of genes in the *fy-1* KO cells, while loss of *fy-2* mainly caused down-regulation of mRNA levels (Figure 5A). Such a bias towards either up- or downregulation was not observed upon KO of *fy-4* or *fy-5* (Figure S5A). In order to analyze the datasets further, the respective mRNAs were subdivided into different groups, according to their occurrence in the different KO cell lines and the direction of regulation (Figure S5B, C). Subsequently, these groups were probed for an enrichment of gene ontology (GO) terms, kegg or reactome pathways. Firstly, we were interested in a common phenotype of the FERRY complex and focused on the groups of mRNAs that were affected by the KO of all four FERRY subunits. Among the downregulated mRNAs the most prominent group was nucleosomal mRNAs. Components of the mitotic spindle and cell junction proteins were found to be upregulated. Further subdivision of the transcriptomic changes allowed us to assess FERRY component-specific phenotypes. Among the mRNAs downregulated in the *fy-2* and *fy-4* KO cell lines, mRNAs of ribosomal proteins and nuclear-encoded mRNAs for mitochondrial protein complexes were enriched. Also, the loss of Fy-2 alone decreased the abundance of the transcripts connected to the ribosome. These data indicate that the loss of FERRY components has a broad influence on the cellular transcriptome, affecting different pathways and cellular processes. The differential effect of the KO of the different subunits of the complex might also reflect additional roles of these components beyond their role in the complex itself. The downregulation of nuclear-encoded mRNAs of mitochondrial proteins and components of the translation machinery in the *fy-2* and *fy-4* KO cell lines, support the association of such mRNAs with the FERRY complex.

**Figure 5:**
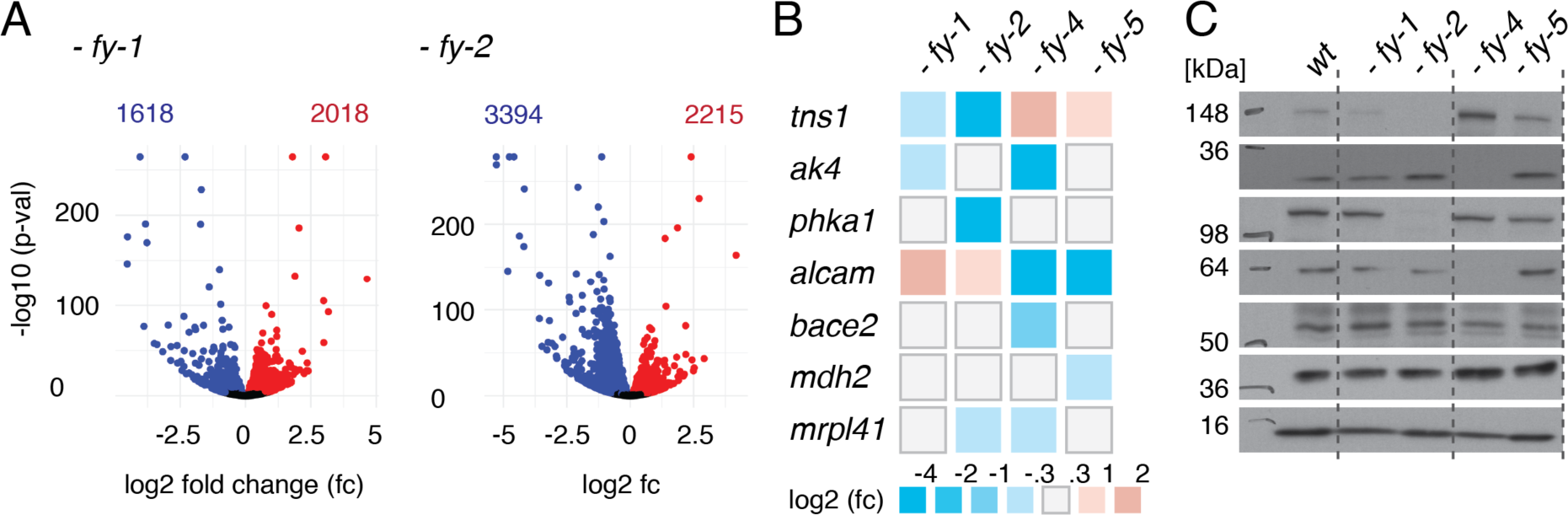
**A)** Volcano plots of the transcriptomic changes in the *fy-1* and *fy-2* KO cell lines compared to *wt*. (blue: down; red: up) **B)** Heatmap of the levels of exemplary mRNAs. **C)** Western blot analysis of the proteins encoded by the mRNAs shown in B.

These finding directly raise the question whether the observed alterations in mRNA levels also translate into changes in protein levels. Therefore, we assessed the levels of proteins translated from mRNAs that are strongly affected by loss of FERRY components (Figure 5B). Western blot analysis showed that changes in mRNA levels can be reflected by strong changes in protein levels, and even manifest in loss of certain proteins, as observed for Tns1 and Phka1 in the *fy- 2* KO and Ak4 and Alcam in the *fy-4* KO cell line (Figure 5C). However, not all changes observed on the transcriptome level were mirrored by changes of protein levels, which can be explained by the wide range of cellular mechanisms that regulate protein levels.

In summary, the FERRY complex not only influences mRNA localization, but also affects the abundance of certain mRNAs which also impacts the respective protein levels. Some of the changes might be a direct consequence of aberrant localization, such as the downregulation of mRNAs for mitochondrial proteins in *fy-2* and *fy-4* KO cell lines. Other effects such as the upregulation of cell junction mRNAs might constitute secondary or compensatory effects of the KO. While aberrant localization of mRNAs in HeLa cells might be compensated by diffusion, such changes likely become detrimental in neurons given their elongated morphology.

### The FERRY complex localizes to axons as well as to the somatodendritic region

Mutations in FERRY subunits have a major impact on brain development and function. Therefore, we assessed the localization of the FERRY complex in primary rat hippocampal neurons. To determine its distribution, we compared the FERRY localization with respect to EEA1 and Rabankyrin-5. EEA1 and Rabankyrin-5 differ in their localization in neurons, as EEA1 is restricted to the somatodendritic region (Wilson et al., 2000), while Rabankyrin-5 is also found in axons (Goto-Silva et al., 2019). Again, we observed a punctate pattern of fluorescent foci dispersed across the neuron for Fy-2 (Figure 6A, overview) and the fluorescent signal strongly co-localized with the endosomal markers EEA1 and Rabankyrin-5. We observed many triple positive (Fy-2, EEA1, Rabankyrin-5) endosomes (Figure 6A, details, white arrowheads), but also fluorescent foci that were only positive for Fy-2 and Rabankyrin- 5, mainly in thin structures devoid of EEA1 (Figure 6A, blue, yellow arrowheads). These results suggest that the FERRY complex is present in both somatodendritic region and axons.

**Figure 6:**
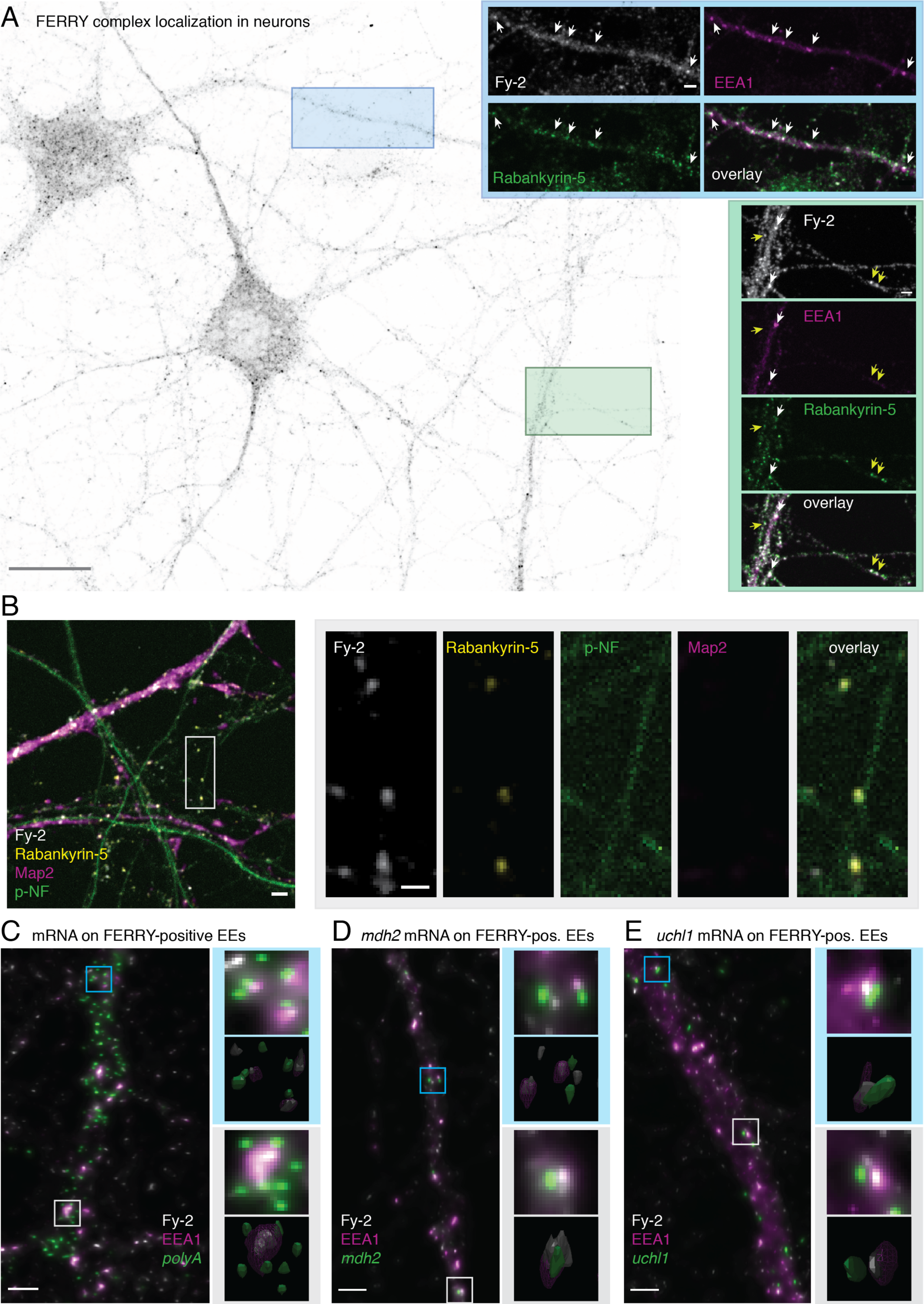
**A)** Localization of the FERRY complex in neurons. Overview image: (Scale bar: 20 μm). The boxed regions (blue and green) are highlighted and shown with additional markers (EEA1, Rabankyrin-5) (Scale bar: 2 μm). White arrowheads indicate co-localization of Fy-2, EEA1 and Rabankyrin-5, yellow arrowheads co-localization of Fy-2 and Rabankyrin-5. **B)** Primary rat hippocampal neurons were stained for Fy-2, Rabankyrin- 5, Map2 and a phosphorylated neurofilament (pNF) (Scale bar: 2 μm). The boxed region is highlighted on the right (Scale bar: 1 μm). **C)-E)** Hippocampal neurons stained for Fy-2, EEA1 and *polyA,* or *mdh2* or *uchl1*. Magnifications and a 3D representation of the indicated regions (grey, blue) are given on the right. (Scale bar: 2 μm).

In order to validate this hypothesis, we performed immunofluorescence against Map2 and the phosphorylated neurofilament-1 (pNF) as markers of the somatodendritic region and axons, respectively (Figure 6B, overview). As our previous experiments suggested, we observed Fy- 2 and Rabankyrin-5 positive EEs in thin structures positive for the axonal marker pNF (Figure 6B, box). In summary, the FERRY complex resides on EEs distributed across the neuronal soma, dendrites and axons, raising the question about possible mRNA localization on these endosomes.

### The FERRY complex co-localizes with mRNA on EEs in neurons

To investigate whether FERRY-positive EEs also carry mRNA in neurons, we visualized the total pool of mRNAs using a *polyA* probe and focused on imaging dendrites and axons, as the cell body has a high protein and mRNA density. While the mRNA density in major dendrites is still high, it decreases in thinner processes and forms clusters at nodes. Overall, we observed that 6.1% of mRNA foci co-localize with the FERRY complex (Figure 6C). Often, these structures also co-localize with EEA1, suggesting that mRNAs are located on EEs (Figure 6C, light blue box). In other cases, a larger endosome is surrounded by several mRNA foci, with the fluorescent signals in close proximity rather than co-localizing (Figure 6C, white box). Deconvolution allowed us to attain a resolution of ∼150 nm in the XY-plane. However, z-resolution of confocal microscope is >500nm. This means that, under normal confocal conditions, fluorescent signals in close proximity to each other would be partially co-localized and within a distance of 250 nm, which is the expected range of a FERRY-mediated attachment of mRNA to the EE, taking into account the antibody labelling and the fluorescent *in situ* hybridization (FISH) (Figure S4B). In case of dense fluorescent signals, similar to those of *polyA*, a substantial proportion of apparent colocalization might result from random co- localization. Therefore, we estimated the random co-localization (Kalaidzidis et al., 2015) (Figure S6A, left).). The results of the analysis indicate that the co-localization of Fy-2 and mRNA is significantly higher than random, indicating the attachment of mRNAs to the EE (Figure S6A, right).

We next tested the co-localization of the FERRY complex with specific transcripts in neurons choosing the *mdh2* and *uchl1* mRNAs based on the initial mRNA-binding screen (Figure 2E) and the co-localization experiments in HeLa cells (Figure 4E). Visualizing individual mRNAs, we obtained much less fluorescent signal per cell (Figure 6D, E). However, we still observed fluorescent signals partially overlapping or in close proximity below 250 nm (Figure 6D, E boxes). Quantifying the number of events, we found 13.2% of *mdh2* transcripts and 10.3% of *uchl1* mRNAs in close proximity or partially overlapping with the FERRY complex.

### mRNA-loaded FERRY-positive endosomes colocalize with mitochondria

The interaction between the FERRY complex and different transcripts encoding mitochondrial proteins suggests that FERRY-positive EEs loaded with mRNAs might be observed in the proximity to, or on, mitochondria for localized translation. To examine this, we additionally stained neurons with TOM70 as a marker for mitochondria. When visualizing the *polyA* mRNA population, we regularly found co-localization of the FERRY complex with mRNA on mitochondria (Figure 7A). We also assessed the co-localization of the FERRY complex with the *mdh2* mRNA and mitochondria (Figure 7B). Even though these events were infrequent, we observed examples where the fluorescence signal of the FERRY complex, the *mdh2* mRNA and mitochondria were in close proximity (Figure 7B, blue box) or even co-localizing (Figure 7B, grey box). Despite the abundance of mitochondria, the degree of co-localization of mRNAs, FERRY and mitochondria are above the expected value for random co-localization, indicating the detection of biologically meaningful events (Figure S6B). These findings support the notion that the FERRY complex is involved in the localization and the distribution of specific mRNAs, such as transcripts encoding mitochondrial proteins (e. g. *mdh2* mRNA), most probably by mediating their endosomal transport (Figure 7C).

**Figure 7:**
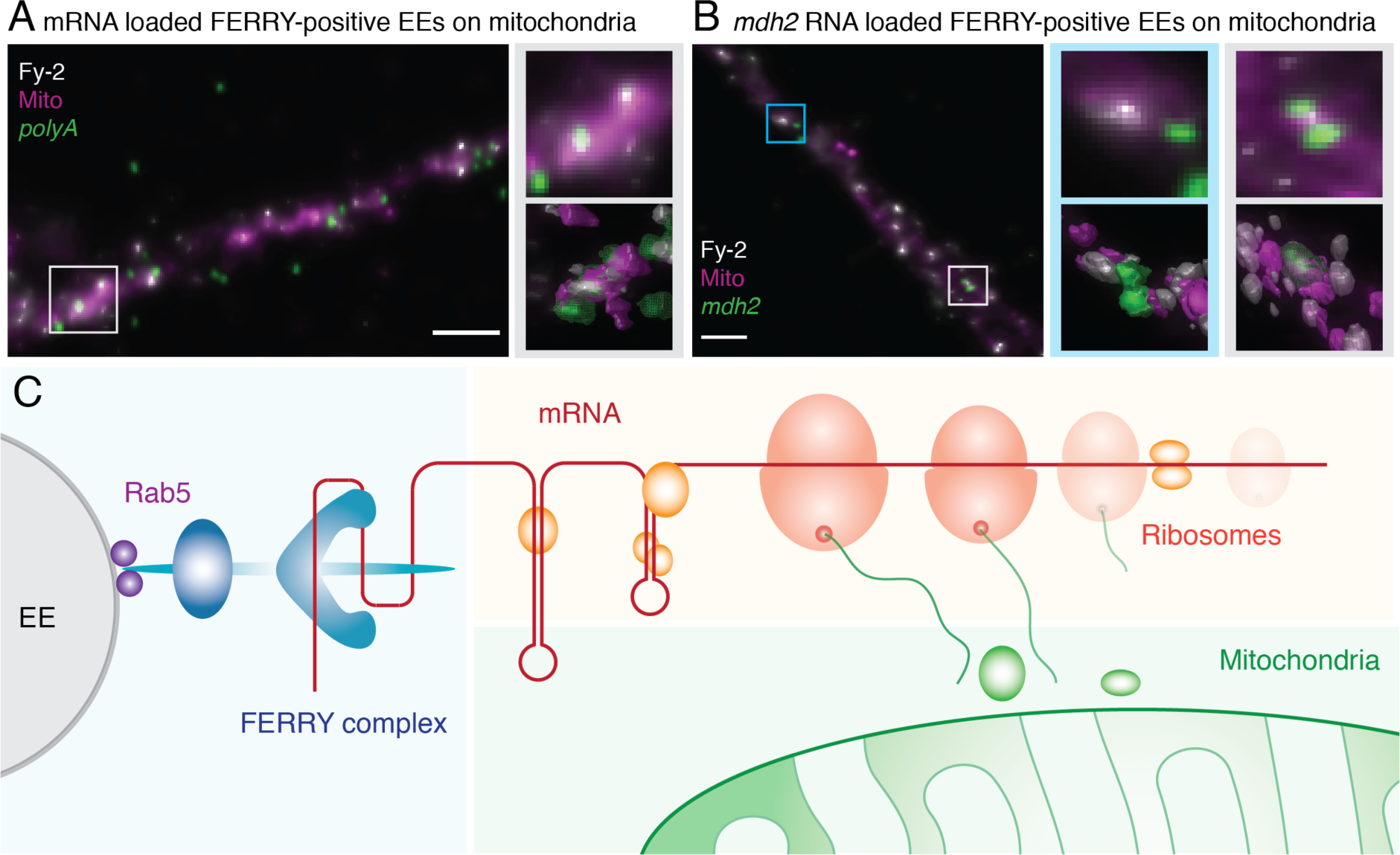
**A)** and **B)** Hippocampal neurons stained for Fy-2, TOM70 and *polyA,* or *mdh2*. Magnifications and a 3D representation of the indicated regions (grey, blue) are given on the right. (Scale bar: 2 μm). **C)** Scheme of the cellular role of the FERRY complex.

## Discussion

### A novel link between the endosomal system and the translation machinery

In this study, we identified and characterized a novel Rab5 effector complex, named FERRY, composed of five subunits, Fy-1 to Fy-5, which interacts with activated Rab5 and is predominantly located on EEs. Furthermore, FERRY directly interacts with mRNAs and in cells, is able to recruit mRNAs onto EEs, enabling the cell to exploit the full logistic capacity of the endosomal system to organize mRNA transport and distribution (Figure 7C). Unlike endocytosed cargo molecules that are inside the endosomal lumen, the RNA is bound and transported on the outside. The FERRY complex therefore couples two vital cellular functions, gene expression and vesicular transport.

### FERRY is a novel type of RNA-binding protein complex

Although the FERRY complex does not contain known RNA-binding motifs, it directly interacts with mRNAs *in vitro* and in cells. Structural studies identified the main mRNA-binding interface as a coiled-coil region at the N-terminus of Fy-2 with additional involvement of several other FERRY components (Quentin et al., 2021). These data indicate that the FERRY complex may also define a novel class of RBPs. The FERRY-RNA interaction is characterized by a large and complex interface on the protein but also on the RNA side. Furthermore, coiled-coil regions have not yet been identified as RNA-binding motifs and thus offer the possibility for further discoveries. Despite the large RNA-binding interface on the FERRY complex, we only detected moderate binding affinities to RNAs *in vitro*. These findings point towards additional layers of regulation in the FERRY-RNA interaction *in vivo*, which could be post-translational modifications, *e.g.* phosphorylation, structural or conformational features of the FERRY complex, or the mRNAs that might be provided by yet unknown factors. Our attempts to subdivide RNAs often led to decreased binding of the individual fragments. This further indicates a similarly complex interface on the RNA that may be composed of multiple distinct motifs distributed along the length of the RNA. Rather than showing a preference for single- or double-stranded mRNA, the FERRY complex might interact with distinct structural elements of certain RNA folds. Taken together, the FERRY complex exhibits novel RNA-binding features, and provides a new model system to obtain deeper insights in RNA-protein interactions in future studies.

### mRNA transport on endosomes

Recent studies have highlighted the vital role of different endosomal compartments for mRNA transport and localization (Cioni et al., 2019; Liao et al., 2019; Popovic et al., 2020). The coupling between the endosomal system and the translation machinery raises the question as to which transcripts bind to endosomes, how many mRNA binding sites can endosomes offer and whether these are provided by different RBPs. The observation of several, up to four mRNA foci, on a single endosome (Figure 5C) suggests that endosomes may be able to accommodate multiple mRNAs. However, it does not answer the question whether these originate from the same RBP or from different mRNA attachment systems. The presence of multiple different physical contacts between endosomes and mRNA is supported by a recent study, showing that transcripts can interact with EEs in a translation-dependent or -independent fashion, pointing towards different mechanisms (Popovic et al., 2020). However, the molecular mechanisms of these binding modes remain to be elucidated.

### Connection between mRNA localization and neurodegeneration

Genetic disruption of the FERRY complex causes severe neurological defects in human patients, especially when Fy-1 and Fy-2 are affected. Most cases report a biallelic frame shift mutation in *fy-1* or *fy-2*, which causes a C-terminal truncation of the respective protein at specific positions. These truncations of either of the two large FERRY subunits has severe impact on the FERRY complex on a molecular level. Already a C-terminal truncation of the last 84 amino acids of Fy-2 leads to intellectual disability and brain abnormalities (Suleiman et al., 2018). On a molecular level, this truncation is sufficient to prevent the interactions between Fy-2 and Fy-1/Fy-3 as well as those with Rab5, hence disrupting the structural integrity of the complex and impairing its proper sub-cellular localization (Quentin et al., 2021). The clinically relevant C-terminal truncations of Fy-1 often affect different domains of the protein including the TBC domain, which is the most conserved part of Fy-1. Additionally, the loss of the TBC Rab GTPase activating domain, might have severe consequences for endocytic trafficking. In summary, the reported mutations have a strong impact on the integrity of the FERRY complex on a molecular level and are therefore likely to induce substantial mis- localization of mRNAs. Our findings confirm that the localization of a large variety of transcripts is affected in FERRY subunit defective cells, making it difficult to identify the cellular pathways that lead to the reported symptoms upon disruption. Further studies are needed to disentangle the mechanisms by which mRNA mis-localization leads to systemic brain damage.

### Limitations of the Study

Our experiments show that the FERRY complex directly interacts with mRNA *in vitro* and in cells. However, the observed moderate binding efficacies *in vitro* seem to contradict the extensive mRNA binding interface on the FERRY complex. This interface comprises the N-terminal coiled-coil of Fy-2 as main contact site and several other contacts with the C-terminus of Fy-2, with Fy-1, Fy-3 and Fy-5 (Quentin et al., 2021). The first limitation lies in the determination of reliable *in vitro* binding constants which is hindered by the structural complexity of a five-subunit complex and mRNA, structural heterogeneity including conformations that are unable to engage in the interaction.

A possible explanation for the weak *in vitro* affinities would be additional factors that interact with FERRY and RNA, *e.g.* RBPs. The GST-FERRY interactor screen did not yield a potential interactor which might be able to fulfil this function. However, we acknowledge that other cellular mechanisms may support the FERRY-RNA interaction *in vivo*. Such mechanisms might involve the transient interaction with proteins that act as loading or unloading platforms for mRNA onto FERRY *e.g.* by modifying the conformation of either FERRY or the mRNA. It is also conceivable that post-translational modifications of FERRY, *e.g.* phosphorylation, modifications of the mRNA, *e.g*. methylation, interaction of the FERRY complex with endosomal lipids or yet unknown mechanisms might be involved in regulating such interaction. Given the number of different mRNAs that are produced in the cell, an intricate regulation of mRNA binding seems more likely than a purely affinity driven FERRY-mRNA interaction.

We also observed that the FERRY complex seems to show different binding efficacies for different mRNAs *in vitro*, which suggests that the FERRY complex is able to provide mRNA specificity (Quentin et al., 2021). However, these observations are limited to *in vitro* experiments and further mechanisms (see above) might contribute to mRNA specificity or mRNA recognition in the cell.

Overall, this study provides a novel molecular player that, due to its evolutionary conservation and requirement for organism physiology, plays an important role in the intracellular localization and translational control of mRNAs exploiting the early endosomes as transport system. The identification of the FEERY complex raises a number of questions that need to be addressed in *ad hoc* structure and function studies. Cells where spatial localization of mRNAs is rate limiting, such as neurons or fungi, are systems of choice to address such questions and test predictions of FERRY complex function.

## Acknowledgements

Firstly, we thank I. Bartnik for excellent technical support and P. Hackert for support with the RNA-IP after crosslinking. We also acknowledge S. Raunser and D. Quentin for valuable feedback regarding the manuscript and the members of the cluster of excellence “Physics of Life” (Deutsche Forschungsgemeinschaft under Germany’s Excellence Strategy—EXC-2068– 390729961—Cluster of Excellence Physics of Life of Technische Universität Dresden) for stimulating discussion. Especially, we would like to thank the following Services and Facilities of the MPI-CBG for their support: The antibody Facility, the light microscopy facility, the mass spectrometry facility, the genome engineering facility and protein expression and purification facility. We also thank the DRESDEN-concept Genome Center (DcGC at CMBC at the TU Dresden) supported by DFG (INST 269/768-1) for technical support. Furthermore, we would like to thank Refeyn Ltd (Oxford, UK) for the use of their Mass Photometer. We also thank the Centre for Information Services and High Performance Computing (ZIH) of the TU Dresden for the generous provision of computing power. This research was financially supported by the Deutsche Forschungsgemeinschaft (DFG, German Research Foundation) - Project Number 112927078 - TRR 83 to M.Z., SFB1190 P04 to K.E.B. and SFB1190, P08/14 to M.T.B., and the Max Planck Society. Open access funding was by the Max Planck Society. J.S.S. was funded by the Deutsche Forschungsgemeinschaft (DFG, German Research Foundation) - Project Number 112927078 - TRR 83.

## Author contributions

Conceptualization, J.S.S. and M.Z.; Software, L.H.; Formal Analysis, J.S.S., C.L., L.H., Y.K., A.T.-P. and M.Z.; Investigation, J.S.S., S.t.D., S.C., J.D.G., S.S., R.S., A.G., K.E.B., M.T.B. and M.Z.; Data Curation, L.H.; Writing – Original Draft, J.S.S. and M.Z.; Writing – Review & Editing, all authors.; Visualization, J.S.S., C.L. and L.H.; Supervision, J.S.S. and M.Z.; Project Administration, J.S.S. and M.Z.; Funding Acquisition, E.M.S and M.Z.

## Data availability

RNA Sequencing (RNA-Seq) data and the respective scripts for the analysis of the RNA-Seq and proteomics data are available in a public repository (https://dx.doi.org/21.11101/0000-0007-EEE3-D).

## Material and Methods

### Molecular Cloning

Human *fy-1* (Tbck, ENSG00000145348, Q8TEA7), *fy-2* (Ppp1r21, ENSG00000162869, Q6ZMI0), *fy-3* (C12orf4, ENSG00000047621, Q9NQ89), *fy-4* (Cryzl1, ENSG00000205758, O95825), *fy-5* (Gatd1, ENSG00000177225, Q8NB37) and *rab5a* (ENSG00000144566, P20339), were amplified by polymerase chain reaction (PCR) using Q5 High-Fidelity DNA polymerase (NEB) and digested using NotI, NcoI, AscI, XhoI, PciI (NEB) according to the manufacturer’s protocol. *fy-5* was cloned into a pET based bacterial expression vector as an N-terminally hexahistidine (His6) tagged variant without cleavage site. *fy-4* was cloned into an expression vector for expression in SF9 cells also carrying a non-cleavable N-terminal His6 tag. *fy-1*, *fy-2* and *fy-3* were cloned into a multi gene construct based on a pBLA vector. For the purification of the FERRY complex *fy-1* carried a cleavable N-terminal His6 tag, the other 2 genes were untagged. To obtain GST-FERRY, *fy-2* carried a cleavable Glutathione-S-transferase (GST) tag, while *fy-1* and *fy-3* remained untagged. *rab5* was used as GST fusion variant in the bacterial expression vectors pGAT2 for GST pulldown assays and pGEX-6P-3 for electrophoretic mobility shift assays (EMSAs). Plasmids and primers used in this study are listed in the resources table (Table S3).

### Virus production and insect cell expression

SF9 cells growing in ESF921 media (Expression Systems) were co-transfected with linearized viral genome and the expression plasmid, and selected for high infectivity. P1 and P2 viruses were generated according to the manufacturer’s protocol. Best viruses were used to infect SF9 cells at 10^6^ cells/ml at 1% vol/vol and routinely harvested after around 48 hours at about 1.5x10^6^ cells/ml. The pellet was suspended in lysis buffer (20 mM HEPES (pH 7.5), 250 mM NaCl, 20 mM KCl, 20 mM MgCl2 and 40 mM imidazole) or SEC buffer (20mM HEPES, pH 7.5, 250mM NaCl, 20mM KCl, 20mM MgCl2) supplemented with a protease inhibitor cocktail, flash frozen in liquid nitrogen and stored at -80 degrees.

### Protein purification

#### Fy-5 and GST-Rab5

For expression of Fy-5 and GST-Rab5, *E. coli* BL21 (DE3) (company) were grown in LB medium under autoinduction conditions using D-(+)-lactose monohydrate at 1.75% (w/v), supplemented with the respective antibiotic (50 μg/ml kanamycin or100 μg/ml ampicillin) at 30 °C under constant shaking (165 rpm). Bacteria were harvested by centrifugation (4000 x g, 20 min, 4 °C), suspended in lysis buffer and subsequently lysed or stored at -80 °C. After sonication the lysate was clarified by centrifugation (22 500 rpm/61 236 x *g*, 20 min, 4 °C) and applied to a HisTrap FF column (GE Healthcare) equilibrated with 10 column volumes (CV) of lysis buffer. After extensive washing with lysis buffer, the proteins were eluted in 10-13 ml elution buffer (20 mM HEPES (pH 7.5), 250 mM NaCl, 20 mM KCl, 20 mM MgCl2 and 500 mM imidazole). Elution fractions containing protein were concentrated using Amicon Ultracel-10K/ Ultracel-30K (Millipore) centrifuge filters and subsequently applied to size exclusion chromatography (SEC) using a Superdex 200 column (HiLoad 16/600 Superdex 200 pg, GE Healthcare) equilibrated in SEC buffer. Fractions were analysed using SDS-PAGE. Protein containing fractions were pooled and concentrated to fit experimental requirements. Protein concentrations were determined by spectrophotometer (NanoDrop Lite, Thermo Scientific).

#### Fy-4

For expression of Fy-4, insect cell suspensions were lysed using sonication, the lysate subsequently clarified by centrifugation (22 500 rpm/61 236 x *g*, 20 min, 4 °C), filtrated using Millex® HV membrane filter units with a pore size of 0.45 μm (Merck Millipore) and applied to a HisTrap FF column (GE Healthcare) equilibrated with 10 CV of lysis buffer. After washing with lysis buffer, the protein was eluted in 10-13 ml elution buffer and concentrated with a centrifuge filter (Amicon Ultracel-30K, Millipore). Thereafter, the protein was applied to SEC using a Superdex 200 column (HiLoad 16/600 Superdex 200 pg, GE Healthcare) equilibrated in SEC buffer. The fractions were analysed by SDS-PAGE. Protein containing fractions were pooled and concentrated according to experimental requirements. The protein concentration was determined by spectrophotometer (NanoDrop Lite, Thermo Scientific).

#### FERRY complex

SF9 cell pellets prior infected with a virus containing Fy-1, Fy-2 and Fy-3 were melted and immediately supplemented with an excess of purified Fy-4 and Fy-5 before lysis. Subsequently, the cells were lysed using a Microfluidizer (LM20, Microfluidics). The lysate was clarified by centrifugation (22 500 rpm/61 236 x *g*, 20 min, 4 °C) and filtrated using membrane filters with a pore size of 0.45 μm (Millex® HV membrane filter units, Merck Millipore). The clarified lysate was supplemented with Ni-NTA agarose (1.3 ml resin/ 1 l insect cell pellet, Qiagen) and incubated for 30 mins at 4 °C on a rotating wheel. Subsequently, the resin was transferred into gravity flow chromatography columns (Poly-Prep® Chromatography Column, Bio-Rad) and washed 3 times with i) 8 CV lysis buffer, ii) 8 CV wash buffer (20 mM HEPES, pH 7.5, 250 mM NaCl, 20 mM KCl, 20 mM MgCl2 and 80 mM imidazole), and iii) 8 CV lysis buffer. The protein was eluted in 1 ml fractions with elution buffer and protein containing fractions were applied to SEC without further concentration, using either a Superdex 200 (HiLoad 16/600 Superdex 200 pg, GE Healthcare) or a Superose 6 increase (Superose 6 Increase 10/300 GL, GE Healthcare) which were equilibrated in SEC buffer. Protein containing fractions were pooled and concentrated according to experimental requirements. Concentration was determined by a spectrophotometer (NanoDrop Lite, Thermo Scientific)

#### GST-FERRY complex

SF9 cell pellets prior infected with a virus containing Fy-1, GST-Fy-2 and Fy-3 were melted and immediately supplemented with an excess of purified Fy-4 and Fy-5. The cells were lysed using a Microfluidizer (LM20, Microfluidics), the lysate was clarified by centrifugation (22 500 rpm/61 236 x *g*, 20 min, 4 °C) and subsequently filtrated using membrane filters with a pore size of 0.45 μm (Millex® HV membrane filter units, Merck Millipore). The clarified lysate was supplemented with Glutathione Sepharose 4B (Cytiva, 2.2 ml resin/1 l insect cell pellet) and incubated for 1.5 h at 4 °C on a rotating wheel. The beads were washed once with 10 ml SEC buffer supplemented with purified Fy-4 and 5 and 2 times with 10 ml SEC buffer. To elute the GST-FERRY complex, the beads were incubated with GSH buffer (20 mM HEPES (pH 7.5), 250 mM NaCl, 20 mM KCl, 20 mM MgCl2, 20 mM GSH) for 1.5 h at 4 °C on a rotating wheel and the beads were removed using filter columns (MoBiTec). The protein complex was concentrated using centrifuge filters (Amicon Ultracel-30K, Millipore) and subjected to SEC using a Superdex 200 column (HiLoad 16/600 Superdex 200 pg, GE Healthcare) equilibrated in SEC buffer. Protein containing fractions were pooled and concentrated according to experimental requirements. Concentration was determined by a spectrophotometer (NanoDrop Lite, Thermo Scientific)

#### Rab5:GTPγS

Expression of Rab5a was performed under autoinduction conditions as described before (Fy-5 and GST-Rab5). Harvested bacterial pellets were resuspended in SEC buffer and lysed using sonication. Glutathione Sepharose 4B (Cytiva) was added to the clarified lysate and incubated for 1.5 h at 4 °C. The resin was washed 3 times with SEC buffer and the protein cleaved off the resin using HRV 3C protease (produced in house) at 4 °C over night on a rotating wheel. Afterwards, the protein was concentrated using Amicon Ultracel-30K (Millipore) centrifuge filters and subsequently applied to SEC using a Superdex 200 column (HiLoad 16/600 Superdex 200 pg, GE Healthcare) equilibrated in SEC buffer. Fractions were analyzed using SDS-PAGE. Protein containing fractions were pooled and concentrated according to experimental requirements. The protein concentration was determined by a spectrophotometer (NanoDrop Lite, Thermo Scientific).

For the nucleotide loading, Rab5 was concentrated using an Amicon Ultracel-30K (Millipore) centrifuge filter, subsequently supplemented with 2.5 mM GTPγS and 250 nM of a GST fusion of the Rab5 GEF domain of Rabex5 (GST-Rabex5-Vps9) and incubated for 1 h on ice. To remove the Rab5 GEF domain, Glutathione Sepharose 4B (Cytiva) was added to the mixture and incubated for 1.5 h at 4 °C. The resin was pelleted by centrifugation (12 000 rpm/ 15 300 x g, 10 min, 4 °C) and the supernatant containing the GTPγS loaded Rab5 was flash frozen and stored at -80 °C. The protein concentration was determined using a BCA assay (Pierce™ BCA Protein Assay Kit, Thermo Scientific).

### GST pulldown assay

5 nmol of purified GST-Rab5 was incubated with 12 µl Glutathione Sepharose 4B (Cytiva) in 100 µl SEC buffer in small filter columns (MoBiTec) for 60 min at 4 °C moderately shaking (700 rpm) in order to saturate the beads with GST protein. Subsequent centrifugation (2500 rpm/660 x *g*, 1 min, 4 °C) removed unbound protein and the resin was washed once with 100 µl SEC buffer. For nucleotide exchange, 1 mM nucleotide (GDP or GTPγS) and 420 nM of GST-Rabex5-Vps9 was added to the columns in 100 µl SEC buffer and incubated for 60 min at 4 °C moderately shaking (700 rpm). After centrifugation (2500 rpm/660 x *g*, 1 min, 4 °C) and subsequent washing with 100 µl SEC buffer, 0.1 nmol FERRY complex was added to the columns in 100 µl SEC buffer and incubated for 20 min at 4 °C on a shaker (700 rpm). Again, unbound protein was removed by centrifugation (2500 rpm/660 x *g*, 1 min, 4 °C) and the columns were washed 3 times with 100 µl SEC buffer. Proteins were eluted with 40 µl of GSH buffer (SEC buffer with 20 mM GSH) for 40 min at 4 °C on a shaker (700 rpm) and analysed by SDS-PAGE and Western blotting.

### Identifying orthologous sequences

We downloaded all eukaryotic reference proteomes from uniport (last accessed: March 2^nd^ 2020) (UniProt, 2019). We used PorthoMCL (Tabari and Su, 2017) to identify orthologous clusters containing human FERRY components (GALD1_HUMAN, QORL1_HUMAN, CL004_HUMAN, PPR21_HUMAN, TBCK_HUMAN). Sequences deviating strongly in length from their human homolog were removed (Table S1). We further distinguished PPR21_HUMAN orthologs between sequences which contain a Fy-4 and a Fy-5 binding site and sequences which do not. For the detection of the presence of the Fy-4 and the Fy-5 binding sites, we aligned all identified Fy-2 sequences. We considered the binding sites present if all of the regions aligned to the PPR21_HUMAN binding regions contained less than 20% gaps (ignoring gapped sites in PPR21_HUMAN).

### Phylogenetic tree estimation

All orthologous clusters were scanned for species which contain at least 80% of identified species with FERRY proteins (custom R script; R 3.6.1; (R Core Team, 2019)). Sequences belonging to FERRY containing species were extracted and aligned using MAFFT with default settings (Rozewicki et al., 2019). Each alignment was trimmed using trimAL (Capella-Gutierrez et al., 2009). The maximum likelihood (ML) tree was estimated using IQTree (Nguyen et al., 2015) whereby each protein was represented as a partition (Chernomor et al., 2016). The Whelan and Goldman matrix (Whelan and Goldman, 2001) with ML optimized amino acid frequencies (WAG+FO) was used as common model for all partitions. Branch support was calculated by IQTree via ultra-fast bootstrapping (UFBoot, 10,000) (Hoang et al., 2018). The consensus tree with the presents/absence information was visualized using the R package ggtree (Version 2.0.4) (Yu et al., 2018; Yu et al., 2017).

### FERRY evolution and ancestral state reconstruction

The identified orthologous genes were used to estimate the ancestral composition of the FERRY complex. The probability for each protein’s presence at each internal node was estimated using Pagel’s algorithm (Pagel, 1994) implemented in the R package ape (Version 5.3) (Paradis and Schliep, 2019).

### Antibody production

Rabbit polyclonal antibodies against Fy-4 were raised in NZW rabbits using standard procedures. 200 ug of recombinant protein emulsified in Complete Freund’s adjuvant was used for immunization. Three boosts were done at 4-week intervals using 200 ug of recombinant protein emulsified in Incomplete Freund’s adjuvant. The final bleed was harvested 10 days after the last boost. Antibodies were affinity-purified on Fy-4 immobilized on a HiTrap NHS- activated HP column (GE Healthcare). Antibodies were eluted using Pierce Gentle Ag/Ab Elution Buffer (ThermoFisher).

Mouse monoclonal antibodies against different components of the FERRY complex were raised in Balb/c mice after subtractive immunization (Sleister and Rao, 2001) with Fy-5. Mice were injected with recombinant Fy-5 in the presence of the immunosuppression drug cyclophosphamide in order to preferentially eliminate Fy-5-reactive B and T lymphocytes. Thereafter the mice were immunized with the entire FERRY complex. Hybridoma were generated using PEG fusions following standard protocols. Clones reacting with individual components of the FERRY complex were selected in a multiplex electrochemiluminescence assay on the MSD platform (Mesoscale Discovery, Rockville, MD). Antibodies were purified from hybridoma supernatant using HiTrap Protein G columns (GE Healthcare).

### Antibody validation

Validation of in-house produced antibodies against components of the FERRY complex for Western blot (WB) were tested against 100 ng, 10 ng and 1 ng of recombinant FERRY complex. Candidates with high sensitivity (detection of 1 ng) and good selectivity (preferably no or no interfering additional signal) were chosen.

Immunofluorescence (IF) validation of the Fy-2 and Fy-4 antibodies raised for this study for was performed using the respective FERRY component KO cell lines (Figure S1F). We subsequently compared the fluorescence signal in wildtype and the KO cell line. For Fy-2 we observed a strong reduction of fluorescence signal in *fy-2* Ko cell line, while the fluorescence of Rabankyrin-5 seems unchanged (Figure S1C, upper panels). Although the WB indicates the disappearance of the Fy-2, we cannot rule out that that there is a small fraction of Fy-2 left. We also tried to generate a KO using a full locus deletion of fy-2, which had a lethal effect on HeLa cells. Thus, we did not obtain any clones. The fluorescence signal for Fy-4 almost completely disappeared in the *fy-4* Ko cell line, while again the Rabankyrin-5 signal seems unchanged (Figure S1C, lower panels).

### Antibodies

The following primary antibodies were used for IF or WB experiments at the concentrations or dilutions indicated: anti-EEA1 (rabbit, polyclonal, laboratory-made, IF 1:1000), anti-Rabankyrin-5 (rat, monoclonal, laboratory-made, IF 1:2000), anti-Map2 (rabbit, polyclonal, Chemicon, IF 1:1000), anti-pNF-H (mouse, monoclonal, Biolegend, IF 1:5000), anti-Fy-1 (rabbit, polyclonal, Sigma Aldrich, HPA039951, WB 1:1000) anti-Fy-2 (mouse, monoclonal, laboratory-made, IF 1:1000, WB 0.5 μg/μl), anti-Fy-3 (rabbit, polyclonal, Sigma Aldrich, HPA037871, WB 1:1000), anti-Fy-4 (rabbit, polyclonal, laboratory-made, IF 1:1000, WB 0.5 μg/μl), anti-Fy-5 (mouse, monoclonal, laboratory-made) WB (0.5 μg/μl), anti-GAPDH (rabbit, monoclonal, Sigma Aldrich, G8795, WB 1:5000), anti-TNS1 (rabbit, polyclonal, Atlas Antibodies WB 1:1000), anti-AK4 (rabbit, polyclonal, Atlas Antibodies, WB 1:1000), anti- PHKA1 (rabbit, polyclonal, Atlas Antibodies, WB 1:1000), anti-Alcam (rabbit, polyclonal, WB 1:1000), anti-BACE2 (rabbit, polyclonal, Atlas Antibodies, WB 1:1000), anti-MDH2 (rabbit, polyclonal, Atlas Antibodies WB 1:500), anti-MRPL41 (rabbit, polyclonal, Atlas Antibodies WB 1:1000), anti-Flag (mouse, monoclonal, Sigma Aldrich, WB 1:10000 or 1:7500), anti-RPL3 (rabbit, polyclonal, Proteintech, WB 1:2000) and anti-RPS3a (rabbit, polyclonal, Proteintech, WB 1:2000).

The following fluorescent secondary antibodies for immunostainings were purchased from Invitrogen and used in a 1:1000 dilution: Goat anti-Rat IgG (H+L) Highly Cross-Adsorbed Secondary Antibody, Alexa Fluor 488, Goat anti-Mouse IgG (H+L) Highly Cross-Adsorbed Secondary Antibody, Alexa Fluor 568, Goat anti-Mouse IgG (H+L) Cross-Adsorbed Secondary Antibody, Alexa Fluor 405, Goat anti-Rabbit IgG (H+L) Cross-Adsorbed Secondary Antibody, Alexa Fluor 647, F(ab’)2-Goat anti-Rabbit IgG (H+L) Cross-Adsorbed Secondary Antibody, Alexa Fluor 647, Goat anti-Mouse IgG (H+L) Cross-Adsorbed Secondary Antibody, Alexa Fluor 488. For WB horseradish peroxidase (HRP) secondary antibodies were supplied from Jackson ImmunoResearch and used at a 1:10 000 dilution.

### HEK 293 lysate preparation

FreeStyle™ 293-F Cells (Thermo Fisher Scientific) were grown in suspension culture in FreeStyle™ 293 Expression Medium (Thermo Fisher Scientific) to density of 4 x 10^6^ cells/ml and harvested by centrifugation (500 x g, 10 min, 20 °C). The cell pellets were suspended in lysate buffer (6 ml/ liter cell culture, 50 mM HEPES (pH 7.5), 100 mM NaCl, 5 mM MgCl2, 1 mM DTT, 0.1% Tween 20), supplemented with a protease inhibitor cocktail and immediately flash frozen in liquid nitrogen. For lysate preparation the pellets were melted, lysed using a microfluidizer (LM20, microfluidics). The lysate was subsequently clarified by a two-step centrifugation (4000 rpm/ 1935 x g, 10 min, 4 °C and 22 500 rpm/ 61 236 x *g*, 25 min, 4 °C), yielding around 15 ml cells lysate per liter cell culture.

### GST-FERRY interactor screens

The GST-FERRY interactor screen was performed at 4 °C in gravity flow filter columns (Poly-Prep® Chromatography Column, Bio-Rad). 500 μl Glutathione Sepharose 4B (GE Healthcare) was added to 0.8 μmol of GST or 7 mg of GST-FERRY complex in 9 ml SEC buffer and incubated for 2.5 h on a rotating wheel. The solution was let run through and the resulting bed of beads was washed 3 x 2 ml SEC buffer. 10 ml of freshly prepared HEK 293 lysate was added to each column and incubated for 1.5 h on a rotating wheel. The lysate was allowed to flow through and another 5 ml of cell lysate was added to each column and also run through the column. The columns were extensively washed with 4 ml lysis buffer and 2 x 5 ml and 2 x 7 ml SEC+ buffer (20 mM HEPES, pH 7.5, 250 mM NaCl, 20 mM KCl, 20 mM MgCl2, 1 mM DTT and 0.1% Tween 20). For the elution of the proteins the columns were incubated with 500 μl of GSH buffer for 40 min on a rotating wheel. The elution fractions were visualized by SDS PAGE and further analysed by mass spectrometry.

To isolate FERRY-associated RNA, from a HEK 293 lysate the GST-FERRY interactor experiment was performed as described with slight modifications. For the elution of the proteins and the associated RNA, RLT buffer from the AllPrep DNA/RNA/miRNA Universal Kit (Qiagen) was supplemented with 1% β-Mercaptoethanol and 20 mM GSH and the pH adjusted to 7.5. The subsequent isolation of nucleic acids was performed using the AllPrep DNA/RNA/miRNA Universal Kit (Qiagen) according to the manufacturer’s protocol. The obtained RNA samples were flash frozen and stored at -80 °C. Prior sequencing, the concentration of the samples was determined by spectrophotometer (NanoDrop Lite, Thermo Scientific) and the samples were analyzed using a 2100 Bioanalyzer (Agilent).

### Mass spectrometry

Samples were separated on SDS PAGE, visualized with Coomassie staining and entire gel lanes cut in 10 pieces each of which was processed individually. Proteins were in-gel reduced by dithiothreitol (DTT), alkylated by iodoacetamide and digested overnight with trypsin (Promega). The resulting peptide mixtures were extracted twice by exchange of 5% formic acid (FA) and acetonitrile, extracts pulled together and dried in a vacuum centrifuge. Peptides were re-suspended in 25µl of 5% FA and 5µl aliquot was analysed by LC-MS/MS on a nanoUPLC system interfaced on-line to a Q Exactive HF Orbitrap mass spectrometer (both Thermo Fischer Scientific). The nanoUPLC was equipped with an Acclaim PepMap100 C18 75 μm i.d. x 20 mm trap column and 75 μm x 50 cm analytical column (3μm/100A, Thermo Fisher Scientific). Peptides were separated using a 80 min linear gradient; solvent A - 0.1% aqueous FA, solvent B - 0.1% FA in acetonitrile. Blank runs were introduced after each sample analysis to minimize carryover. Instrument performance was monitored with QCloud system (Chiva et al., 2018). Data were acquired using a Top 20 approach; precursor m/z range was 350-1600 and dynamic exclusion time was 20 s. The lock-mass function was set on the background ion (Si(CH3)2O)6 at m/z 445.12. Acquired spectra were converted into the .mgf format and merged into a single file for each sample.

Acquired data were processed with the MaxQuant software package (v.1.6.10.43, (Cox and Mann, 2008)) using default setting iBAC options, with Match-Between-Runs (MBR) disabled. Enzyme specificity was trypsin, number of allowed miscleavages – two; variable modification – cysteine carbamidomethyl, propionamide; methionine oxidation; protein N-terminus acetylated.

### Mass photometry

Mass Photometry (MP, iSCAMS) of the FERRY complex was performed on a One^MP^ instrument (Refeyn, Oxford, UK) at room temperature. High precision 24 x 50 mm coverslips (Thorlabs CG15KH) were cleaned with ultrasound, rinsed with isopropanol and water and dried with clean nitrogen gas (Young et al., 2018). 20 µl diluted FERRY complex (43 and 34 nM, in PBS) was spotted into a reusable culture well gasket with 3 mm diameter and 1mm depth (Grace Bio-Labs). MP signals were recorded for 60 s at a suitable concentration in order to detect a sufficient set of target particles (>500). Raw MP data were processed in the DiscoverMP software (Refeyn, Oxford, UK).

### Sucrose density gradient centrifugation to analyze ribosome association

Expression of 2xFlag-PreScission protease cleavage site-His6-Fy2 was induced in stably transfected HEK293 cells by addition of 1 μg/μl tetracycline for 24 h. Cells were treated with 100 μg/ml cycloheximide for 10 min prior to harvesting. Cells were resuspended in Lysis Buffer (20 mM HEPES pH 7.6, 100 mM KCl, 5 mM MgCl2, 0.5% NP-40, 100 μg/ml cycloheximide, 2 mM DTT, 0.625% Triton-X-100, 0.625% deoxycholate supplemented with protease and RNase inhibitors) and lysed on ice for 5 min. Cell debris were pelleted by centrifugation at 10,000 g for 10 min at 4 °C. Extracts were separated on 10-50% sucrose gradients prepared in Lysis Buffer lacking detergents by centrifugation in an SW-40Ti rotor at 35,000 rpm for 2.5 h (Jaafar et al., 2021). Gradients were fractionated and an absorbance profile at 260 nm generated using a BioComp Gradient Master (Aquino et al., 2021). Relevant fractions were pooled and proteins precipitated using 20% trichloroacetic acid. Proteins were separated by SDS-PAGE and analyzed by WB using anti-Flag (Sigma-Aldrich F3165; 1:7500), anti-RPL3 (Proteintech 11005-1-AP; 1:2000) and anti-RPS3a (Proteintech 14123-1-AP; 1:2000) antibodies.

### Library preparation and Sequencing

mRNA was enriched from 100ng DNase treated total RNA using the NEBNext rRNA depletion Kit (human, mouse, rat, NEB) according to the manufacturer’s instructions. Final elution was done in 5 µl nuclease free water. Samples were then directly subjected to the workflow for strand specific RNA-Seq library preparation (NEBNext Ultra II Directional RNA Library Prep, NEB). 0.15 µM NEB Adaptor were used for ligation. Non-ligated adaptors were removed by adding XP beads (Beckmann Coulter) in a ratio of 1:0.9. Dual indexing (GST- FERRY association screen) or unique dual indexing (RNASeq of FERRY component KO cell lines) was done during the following PCR enrichment (12 cycles, 65°C)). After two more XP bead purifications (1:0.9) libraries were quantified using the Fragment Analyzer (Agilent). Libraries were equimolarly pooled before sequencing them with a length of 75 bp in single end mode on an Illumina NextSeq 500 system to a depth of at least 2 x 10^7^ reads (GST-FERRY association screen) or with a length of 2 x 150 bp in paired end mode on an Illumina NovaSeq 600 system to a depth of at least 5 x 10^7^ read pairs (RNASeq of FERRY component KO cell lines).

### Analysis of the mass spectrometry data

From the MaxQuant proteinGroups.txt file only protein groups with at least 1 unique peptide and which were identified in at least two out of three biological replicates in at least one condition were considered for differential abundance analysis using DEP v1.4.0 (Zhang et al., 2018). After variance stabilizing normalization (Huber et al., 2002) of iBAQ intensities, missing values were imputed applying the nearest neighbor averaging imputation method (KNN) to missing at random (MAR) and left-censored imputation using a deterministic minimal value approach (MinDet) to missing not at random (MNAR) protein groups (Gatto et al., 2021). MNARs refer to those protein groups with missing values in all replicates of one of the two conditions while all other missing values are considered as MAR. The application of empirical Bayes statistics on protein group-wise linear models was done using limma (Ritchie et al., 2015) and differentially abundant proteins were identified by applying a log2 fold change threshold of 1 and an adjusted p-value cutoff of 0.05.

### Analysis of the RNA sequencing data

Raw reads were checked for their overall quality using FastQC v0.11.2 (Andrews, 2010). Read mapping to the human genome reference assembly (GRCh38_p13) and genes counts estimation based on Ensembl release v99 (Yates et al., 2020) were done using STAR v2.5.2b (--outFilterMultimapNmax 1 --outSJfilterCountUniqueMin 8 3 3 3 --quantMode GeneCounts; (Dobin et al., 2013) by taking read strandedness into account. Count data were filtered for genes with more than 10 counts in any sample and served as input for differential gene expression analysis using DESeq2 v1.22.1 (Love et al., 2014). An adjusted p-value cutoff of 0.01 was applied to FDRs obtained by using IHW v1.10.1 (Ignatiadis et al., 2016). Results summary in form of a MA plot was done using ggplot2 v3.2.1 (Wickham, 2016) following layout settings from the ggpubr package v0.2.5 (Kassambara, 2020). A gene set enrichment analysis against the MSigDB C5 collection of ontology sets (msigdbr v7.2.1, (Dolgalev, 2020)) was run using fgsea v1.8.0 (Korotkevich et al., 2021) excluding gene sets with less than 15 and more than 500 genes (Subramanian et al., 2005).

### Rab5 affinity chromatography

GST-Rab5 affinity chromatography was carried out as described before (Christoforidis et al., 1999). In summary, GST-Rab5:GDP or GST-Rab5:GTPγS loaded glutathione Sepharose was incubated with bovine brain cytosol, the beads extensively washed and the bound proteins subsequently eluted. The resulting mixture of Rab5 effector proteins was further purified by SEC and anion exchange chromatography. Fractions were analyzed using silver stained SDS PAGE.

### *In vitro* translation binding assay

Binding assays with *in vitro* translated proteins were essentially performed as described (Nielsen et al., 2000). Briefly, [^35^S]-methionine-labelled proteins were transcribed and translated using a TnT™ coupled transcription–translation kit (Promega) according to the manufacturer’s protocol. Resulting proteins were incubated with GST-Rab5:GDP or GST- Rab5:GTPγS loaded Glutathione Sepharose for 2 h at 4 °C. Subsequently, the beads were washed and Rab5-bound proteins were eluted and analyzed by SDS PAGE and fluorography as described (Christoforidis et al., 1999).

### mRNA production and electrophoretic motility shift assays (EMSAs)

mRNA sequences for *mrpl41*, *mdh2*, *uchl1*, *atp5f1b*, *gstp1*, *prdx5*, *cox6b*, *cox8a* and *pigl* comprise the coding region, the 3’ and 5’ untranslated regions (UTRs) and an additional polyA appendix of 50 adenines (Table S3). The mRNAs were produced by *in vitro* transcription using the T7 RiboMAX™ Express Large Scale RNA Production System (Promega) according to the manufacturer’s protocol. Resulting RNA was purified using a Phenol:Chloroform extraction and an isopropanol precipitation as described in the manual of the mMESSAGE mMACHINE™ T7 Transcription kit (Thermo Fisher). In brief, the *in vitro* transcription reactions were quenched with Ammonium acetate stop solution from the mMESSAGE mMACHINE™ T7 Transcription Kit (Thermo Fisher) and supplemented with Phenol:Chloroform:Isoamyl Alcohol 25:24:1 (Sigma Aldrich). The aqueous phase was recovered and RNA precipitated by adding equal amounts of isopropanol. After chilling at - 20 °C for at least 15 min, the precipitated RNA was pelleted by centrifugation (20 800 x g, 15 min, 4 °C), the supernatant removed and the pellet resuspended in RNAse-free water. RNA concentrations were determined by spectrophotometer (NanoDrop Lite, Thermo Scientific)’ and the RNA was stored at - 80 °C until usage.

For direct protein-RNA interaction assays, 10 pmol of FERRY complex was mixed with *in vitro* transcribed mRNA in varying protein/RNA ratios in SEC buffer in a total volume of 20 μl and incubated for 80 min at 37 °C. The samples were analyzed using gel electrophoresis with 1% agarose gels. Gels were always run as duplicates and one gel stained for RNA using SYBR™ Gold Nucleic Acid Gel Stain (Invitrogen) the other stained for proteins with SYPRO Red Protein Gel Stain (Sigma Aldrich). Both dyes were used according to the manufacturers’ protocols.

Direct protein-RNA interaction assays in presence of Rab5:GTPγS were performed, with 15 pmol of *mrpl41* mRNA mixed with 15 pmol FERRY complex and varying amounts of Rab5:GTPγS as indicated in the respective figure in SEC buffer in a total volume of 35 μl. The mixture was incubated for 80 min at 37 °C and the samples were analyzed by ethidium bromide-stained gel electrophoresis using 1% agarose gels.

### RNA immunoprecipitation after UV crosslinking

Stably transfected HEK293 cell lines for the tetracycline inducible expression of 2xFlag- PreScission protease cleavage site-His6-Fy2, Fy2-His6-PreScission protease cleavage site- 2xFlag or the tag alone were generated using the HEK293 Flp-In T-REx system (ThermoFischer Scientific). Expression of the transgenes was induced by addition of 1 μg/μl tetracycline for 24 h, and cells were grown in the presence of 100 μM 4-thiouridine for 9 h before crosslinking with 360 mJ/cm^2^ irradiation at 365 nm (Choudhury et al., 2019; Memet et al., 2017; Sloan et al., 2015). Cells were harvested, resuspended in a buffer containing 50 mM Tris-HCl pH 7.8, 150 mM NaCl, 1.5 mM MgCl2, 0.1% NP40, 5 mM ′3-mercaptoethanol, cOmplete-EDTA-free protease inhibitors and lysed by sonication. RNA-protein complexes were retrieved from the cleared lysate on anti-Flag M2 beads (Sigma Aldrich) and eluted using 3x Flag peptide. Co-purified RNAs were subjected to partial RNase digestions using RNace-It (Agilent Technologies) and complexes were immobilized on Ni-NTA under denaturing conditions (50 mM Tris-HCl pH 7.8, 300 mM NaCl, 10 mM imidazole 6 M guanidium-HCl, 0.1% NP40, 5 mM ′3-mercaptoethanol). Alkaline phosphatase treatment was performed before labelling of the RNA fragment 5’ ends with [^32^P] using T4 PNK. Complexes were eluted from the Ni-NTA using imidazole and precipitated with 20% trichloroacetic acid before separation by denaturing polyacrylamide gel electrophoresis and transfer to a nitrocellulose membrane. Labelled RNAs in the eluate were then detected by autoradiography and proteins were subjected to WB using an anti-Flag antibody (Sigma-Aldrich F3165; 1:10000).

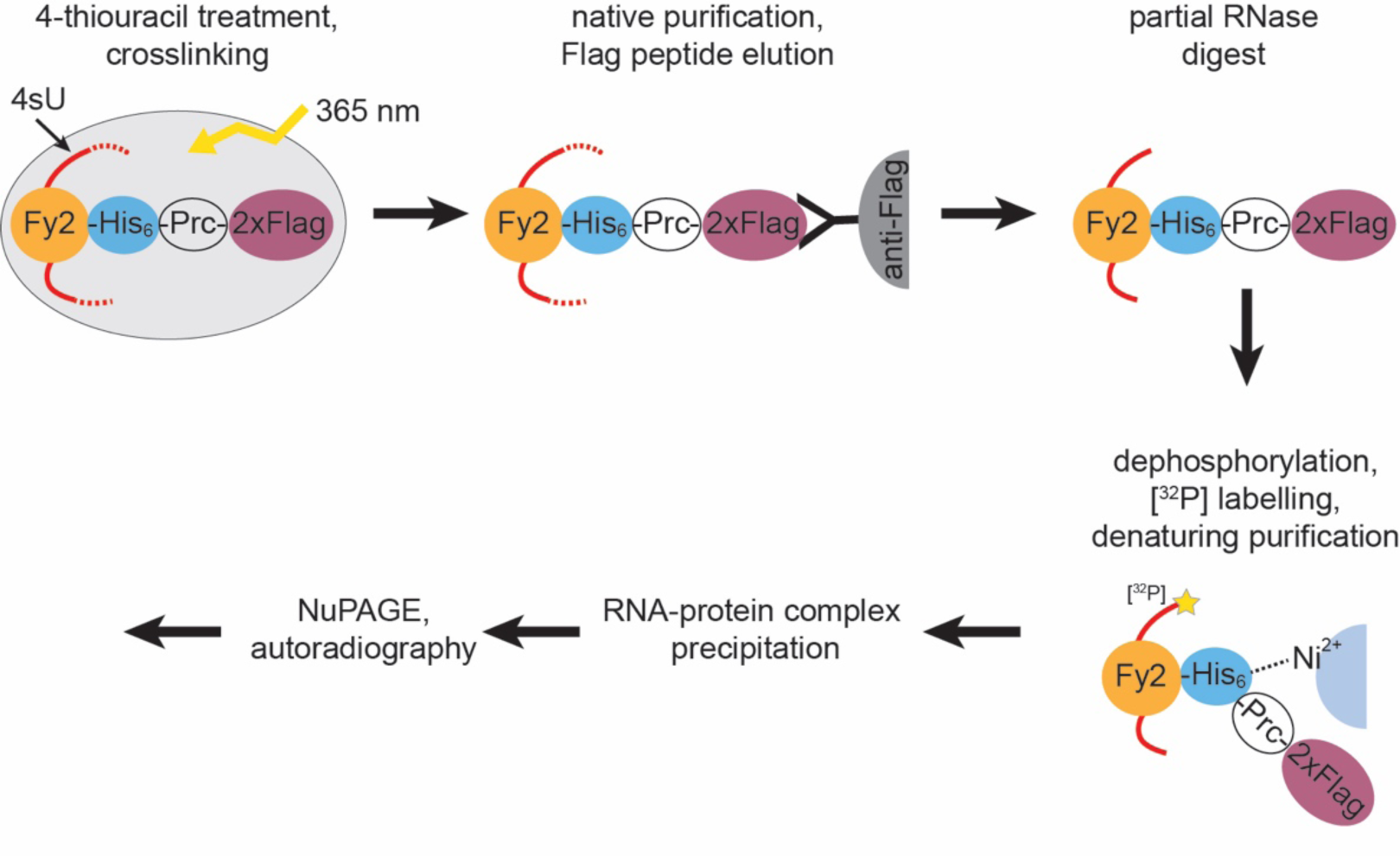

### Generation knockout (KO) cell lines

#### Generation HeLa knockout (KO) cell lines by induced random mutations

To generate gene knockouts in HeLa, we used CRISPR/Cas9 cleavage induced random (NHEJ mediated) mutations using guide RNAs targeted 5’ end of the coding sequence of the genes of interest. We used electroporation of Cas9 protein complexed with crRNA and trRNAs (altR, IDT), using the Neon electroporator device and kits (Invitrogen) with concentrations and electroporation settings as previously described (Spiegel et al., 2019). For list of crRNA protospacers used for each gene, see the resources table (Table S3). The success of the gene disruption was initially assessed by Western blot of single cell derived clones. The disruption of the target alleles was further confirmed by fluorescent PCR and Sanger sequencing of PCR amplicons (For the genotyping primers used and description of the alleles, see the resources table (Table S3).

#### Generation of a *fy-2* KO in HeLa cells by critical exon deletion

In order to generate a *fy-2* knockout in HeLa cells, we deleted exon 6 to 7 (deletion of ca. 1340 bp). Deletion of these two exons generates an out-of-frame transcript with a premature stop codon which leads to a truncated protein of 187 aa. Guide RNAs specific to the *fy-2* locus were selected based on low off-target activity using http://crispor.tefor.net. The guide RNAs were ordered as crRNA from Integrated DNA Technologies (IDT).

HeLa cells were transfected with Cas9 protein (IDT Cat.no. 1081061) complexed with crRNA (IDT, Alt-R®) and tracrRNA (IDT Cat.no. 1072534) using the Neon electroporator device and kits (Invitrogen) with concentrations and electroporation settings as previously described (Spiegel et al., 2019). For a list of crRNA protospacers used for each condition, see the resources table (Table S3). 72 h post-transfection cells were single-cell sorted into 96-well plates. Cell sorting was performed in a BD FACSAria Fusion flow cytometer (Beckton Dickinson). Single-cell clones were genotyped by PCR. Briefly, genomic DNA was extracted using the QuickExtract DNA extraction kit (Epicentre) following the manufacturer’s instructions. PCR was performed using Phusion Flash High-Fidelity PCR Master Mix (ThermoFisher) with gene-specific primers. Amplicons of the deleted alleles were verified by Sanger Sequencing. For the genotyping primers used and description of the alleles, see the resources table (Table S3).

### HeLa cell culture

Hela Kyoto and FERRY subunit knock-out cells were cultured in DMEM media supplemented with 10% FBS Superior (Merck) and 50 μg/ml streptomycin (P/S) (Gibco) at 37°C with 5% CO2. For smFISH studies, cells were seeded into 384 well plates at a density of 3000 cells/well in 50 µl using the drop dispenser (Multidrop, Thermo Fischer Scientific) and cultured for 24h.

### Single molecule fluorescence in situ hybridization (smFISH) and immunostaining

Endosomes and endogenous mRNAs were stained by using the ViewRNA® Cell Plus Assay kit (Invitrogen, 88-19000). The kit consists of 16 solutions that are used to perform an immuno- fluorescence staining followed by a single molecule fluorescence in situ hybridization (smFISH) using the sequential branched-DNA amplification technique. The manufactures protocol for 96 well plates was adapted to a 384 well plate format by down-scaling to 12.5 µl/well for steps containing staining solutions and to 25 µl/well for steps containing washing/fixing solutions (96 well protocol: 50 µl and 100 µl, respectively). For details see the manufactures protocol (https://assets.thermofisher.com/TFS-Assets/LSG/manuals/88-19000.pdf).

In brief, all steps were performed manually using an 8-channel aspirator for removal and automated multi-channel pipettes for addition of liquids. All wash steps following fixation and immunostaining were done 3 times with PBS including RNase inhibitor solution, whereas all wash steps following smFISH were done 5 times with RNA wash buffer solution. Cells were fixed and permeabilized using the provided solutions of the kit. After washing with PBS, cells were incubated with blocking buffer, primary antibody solution (including EEA1 and Fy-2 antibodies at a dilution of 1:2000 and 1:1000, respectively) and secondary solutions (including antibodies against rabbit and mouse IgG labelled with Alexa 488 or Alexa 568 (Alexa 647 for probe HPRT1), respectively, at a dilution of 1:500). After immunostaining cells were fixed and ready for smFISH. Different probes were used to label different mRNAs (Invitrogen, all probes were of type 6 (647nm), except the house-keeping gene HPRT1 (type 1, 546nm); *atp5f1b*: VA6-3168504, *gla*: VA6-3168560, *gstp1*: VA6-3169160, *cox6b*: VA6-3171299, *cox8a*: VA6-3171305, *mdh2*: VA6-3172506, *mrpl41*: VA6-3169863, *mrps35*: VA6-3179781, *psma1*: VA6-3173135, *polyA*: VF6-12675, *rims1*: VA6-3176214 and *hprt1*: VA1-11124).

Cells were incubated for 2h at 40°C with a diluted probe. After washing the cells with RNA wash buffer solution, the protocol was continued the next day with the smFISH branched-DNA amplification technique steps. Subsequently, cells were incubated with pre-amplifier, amplifier and label solution each for 1h at 40°C. Finally, the cells were stored in PBS containing DAPI (1µg/ml) to stain the nuclei and CellMaskBlue (CMB) (0.25µg/ml) to stain the cytoplasm.

### Preparation of hippocampal cultures

Primary rat hippocampal neurons used in this study were obtained and cultured in two different ways. For initial Fy-2 localization experiments, the protocol for culturing hippocampal neurons was adapted from (Goto-Silva et al., 2019) with slight modifications. In brief, neurons were isolated from rat embryos at E17. The rat hippocampi from embryos of either sex were dissected in PBS (25 mM Na-phosphate buffer, pH 7.4, 110 mM NaCl, 1 mM EDTA) and dissociated in digestion solution (100 mg/ml DNAse I and 200 Units Papain in PBS) for 20 min. After two washes of the tissue with plating medium (DMEM containing 10% FCS, 2 mM glutamine, 50 mg/ml penicillin/streptomycin, Invitrogen), it was triturated in plating medium and subsequently cells counted. The neurons were plated on glass cover slips coated with 1 mg/ml poly-L-lysine (Sigma-Aldrich) at a density of 25 000 cells/ml in the presence of a mouse astrocyte feeder layer, derived from the mouse cortex from mice of age P0-P3 of either sex (Kaech and Banker, 2006).

Primary neurons for mRNA localization experiments were obtained and cultured according to the following protocol. Neuronal cultures were prepared from dissociated hippocampi of P0/P1 SD rats as previously described (Cajigas et al., 2012). Hippocampi were collected in Dissociation Medium on ice (DM with 1 mM HEPES, 82 mM Na2SO4, 30 mM K2SO4, 5.8 mM MgCl2, 0.252 mM CaCl2, 20 mM Glucose, 0.001% Phenol Red) and treated with cysteine-activated papain solution in DM (10 ml DM, 3.2 mg Cysteine, 300 µl Papain Sigma P3125, pH readjusted to 7, filtered sterile) two times 15 min at 37°C before several washes with cold DM and Neuronal growth medium (NGM: Neurobasal A supplemented with B27 and Glutamax). Dissociation of the tissue was achieved by trituration through a 10 ml pipette for 10 times. Before counting in a Neubauer chamber, cells were pelleted by centrifugation for 5 min, 67 x g at 4 °C, resuspended in cold NGM and 30 000 cells were seeded in 250 µl NGM on poly-D-Lys coated 14 mm MatTek glass bottom dishes. After attachment of the cells (2-3 h later) 0.7 ml conditioned NGM (80% NGM, 15% glia-conditioned NGM, 5% cortical neuron-conditioned NGM) was added and regular feeding by addition of NGM was performed thereafter. The neurons were kept in an incubator at 37°C in a humidified atmosphere with 5% CO2.

### Animals

The rat pups were used without gender determination. Timed pregnant rats were purchased from either Janvier (RjHan:WI - Wistar rats) or Charles River Laboratories, maintained under food and water ad libitum in a 12h-12h light dark cycle until pups were delivered, pups were sacrificed shortly after birth by decapitation with sharp scissors before dissection of the tissue. The procedures involving animal treatment and care were conducted in conformity with the institutional guidelines that are in compliance with the national and international laws and policies (DIRECTIVE2010/63/EU; German animal welfare law, FELASA guidelines) and approved by and reported to the local governmental supervising authorities (Regierungspräsidium Darmstadt and Landesdirektion Sachsen). The animals were euthanized according to annex 2 of §2 Abs. 2 Tierschutz-Versuchstier-Verordnung.

### Immunostaining of neurons

Immunostaining was performed at room temperature and the plates were subsequently stored at 4 °C if necessary. After adhesion, cells were washed once with PBS and fixed using 3% Paraformaldehyde (PFA) for 15 min. After washing with PBS, residual PFA was quenched using 500 mM Ammonium chloride in PBS for 10 min and the cells were washed 3 times with PBS. For permeabilization the cells were treated with 0.1% Triton X-100 in PBS for 3 min and subsequently washed three times with PBS. After blocking with 10% FBS for 20 min, the cells were incubated with the primary antibody for 2 h. Before and after the application of the secondary antibody for 1 h, the cells were washed 3 times with PBS.

### High sensitivity FISH and immunostaining in neurons

In situ hybridization was performed using the ViewRNA ISH Cell Assay Kit (Thermo Fisher) according to the manufacturer’s protocol with the modifications described previously (Cajigas et al., 2012). Probe sets targeting the respective mRNAs were purchased from Thermo Fisher. In brief, rat hippocampal neuron cultures grown for two weeks on MatTek glass bottom dishes were fixed for 20 min with PBS containing 1mM MgCl2, 0.1 mM CaCl2, 4% Sucrose and 4% PFA, pH 7.4 at room temperature, washed and subsequently permeabilized for 3 min with the provided detergent solution. Gene specific type1 (*uchl1*) and type6 (*mdh2*, *polyA*) probe sets were applied in 1:100 dilution for 3 h at 40°C. After several washes signal amplification steps with PreAmp/Amp and Label Probe reagents coupled to a 550 nm dye were all performed for 1 h at 40°C followed by washes at room temperature after each step. All probe sets and branched DNA reagents were diluted in the provided solutions 1:100. Immunostaining for Fy- 2, endosome and mitochondria markers was performed after completion of the FISH protocol. FISH-stained cells were blocked for 30 min in blocking buffer (BB) at room temperature (BB: PBS with 4% goat serum) and incubated with primary antibodies in BB for 1 h at room temperature. After washing, secondary antibodies in BB were applied for 30 min, cells were washed and nuclei stained by a 3 min incubation with 1 µg/µl DAPI in PBS. Cells were washed in PBS and mounted with Aquapolymount (Polysciences).

### Microscopy

#### automated HeLa imaging

Confocal imaging was performed on an automated spinning disc confocal microscope (Yokogawa CV7000) using a 60x 1.2NA objective. DAPI and CMB was acquired with a laser excitation at 405 nm and an emission band pass filter BP445/45, Alexa 488 with a 488 nm laser and BP525/50 filter, Alexa 568 with a 561 nm laser and BP600/37 filter, Alexa 647 with a 640 nm laser and a BP676/29 filter. 9 fields were acquired per well as a stack with 4 z-planes and 1 µm distance. Each condition was done in duplicate wells and three independent experiments.

#### Spinning disk neuron imaging

Neurons were imaged on a Nikon TiE spinning disk microscope equipped with a 100x/ 1.45NA Plan Apochromat, DIC oil immersion objective, Yokogawa CSU-X1 scan head and a Andor DU-897 back-illuminated CCD detector. Images were acquired with 600 ms exposure, while the laser intensities were adapted to the respective antibodies and requirements. Overview images of almost entire neurons were taken as a set of individual small images (6 x 6 images) with an overlap of 5% and combined using the Fiji implemented Grid/Collection Stitching tool (Preibisch et al., 2009) without overlap computation.

#### confocal neuron imaging

Images were acquired with a LSM780 confocal microscope (Zeiss) equipped with Zen10 software using a 63x/1.46-NA oil objective (alpha Plan Apochromat 63×/1.46 oil DIC M27) and Argon 488, DPSS 561 and HeNe 633 laser lines for excitation in single tracks and a MBS488/561/633 beam splitter. Images were acquired in 12-bit mode as z-stacks and a time series with 4x Zoom, 512px x 512 px resolution and 0.1 µm Tetraspec beads (ThermoFisher) imaged under the same conditions. The laser power and detector gain in each channel was set to cover the full dynamic range but avoid saturated pixels.

### Image analysis

#### HeLa cell images

Microscopy images for the localization of Fy-2, EEA1 and different mRNAs in HeLa cells were processed using the stand-alone freely available software MotionTracking (MT) (http://motiontracking.mpi-cbg.de). Images of were imported into MT and subsequently corrected for the chromatic shift of individual channels based on images of Tetraspec beads. For quantification, fluorescent foci of EEA1 and mRNA were detected using automated object detection and the co-localization was calculated based on 0.35 overlap threshold (Collinet et al., 2010; Kalaidzidis et al., 2015). Colocalization markers on endosomes with and without Bayesian correction for random colocalization was performed using MT as is described in (Kalaidzidis et al., 2015).

#### Neuron images

Microscopy images for the localization of Fy-2, EEA1, mRNA and mitochondria in neurons were also processed with MT. Image sequences of fixed neurons were imported into MT and drift corrected and deconvoluted by algorithms implemented in MT. In a last step, images were corrected for the chromatic shift of individual channels based on images of Tetraspec beads before and after the imaging. Motion Tracking implemented object detection was used to determine the mRNA foci while subsequent image analysis and quantification was performed by visual inspection. Given the possible distance between the fluorescence signals of EEA1 and mRNA or Fy-2 and mRNA (Figure S4A), automated object detection followed by a co- localization analysis was not suitable for this purpose.

#### Multiple source localization microscopy (Musical)

Samples were prepared in fresh STORM buffer as described in (Franke et al., 2019). Image stack acquisition was performed with a Spinning Disc, Andor- Nikon TiE inverted stand microscope equipped with a spinning disc scan head (CSU-X1; Yokogawa), a fast piezo objective z-positioner (Physik Instrumente), a back-illuminated EMC CD camera (iXon EM+ DU-897 BV; Andor), a Nikon Apo 100x 1.45 Oil DIC objective and a OptoVar 1.5 lens (pixel size in x-y plane is 70.1nm). Samples were z-scanned for 2.5μm with 0.25μm steps. At each z- position for 3 channels (488, 561 and 647) 50 snap-shot images were acquired and 405 nm laser was used to re-activate fluorophores before moving to the next z-position. The multiple fluorophore localization was performed by algorithm described in (Agarwal and Machan, 2016) as it implemented in MT.

### Western blotting

Cells were collected from a 10 cm cell culture dish, washed with cold PBS and subsequently lysed in PBS supplemented with 1% Triton X-100. HeLa cell lysates were clarified by centrifugation (14 000 rpm/ 20 800 x *g*, 15 min, 4 °C) and the concentration determined using a BCA assay (Pierce™ BCA Protein Assay Kit, Thermo Scientific). After running an SDS PAGE (12%), the gel was subsequently transferred onto a nitrocellulose membrane (Amersham). Blots were washed with PBST (PBS supplemented with 0.1% Tween 20) and then incubated with WB blocking buffer (5% non-fat milk powder in PBST) over night at 4 °C. After washing with PBST blots were then incubated with the primary antibodies (anti-Fy-1 to anti-Fy-5 and anti-GAPDH as a loading control) at the dilutions indicated earlier for 1 h at room temperature. After washing the secondary HRP antibody was applied to the blot for 1 h at room temperature. All antibodies were added in PBST with 5% milk. The blots were developed using ECL™ Western Blotting Reagents (Cytiva) on respective films (Amersham) in a Kodak X-OMAT 200 Processor.

