## Supplementary Figures for "The novel Rab5 effector FERRY links early endosomes with the translation machinery"

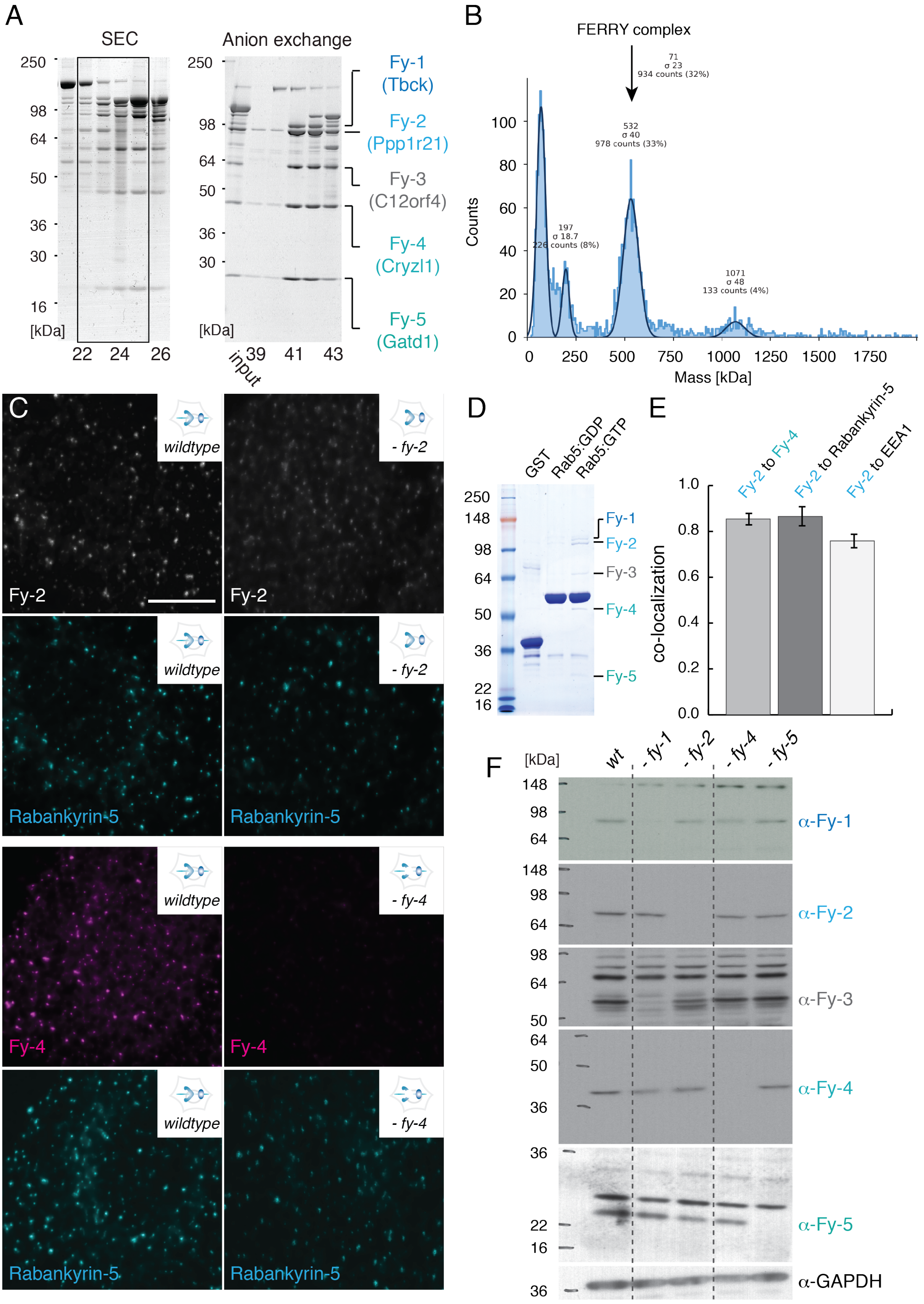


**Figure S1**: **A)** The Rab5 effectors obtained by Rab5 affinity chromatography were fractionated by two chromatographic techniques, i) size exclusion chromatography (SEC, left) and ii) anion exchange chromatography (right). The fractions were analyzed by SDS-PAGE and Coomassie staining. The five proteins of the FERRY complex co-eluted in fractions 22 to 25 from SEC (left gel). Fractions 22 to 25 were combined and subjected to an anion exchange chromatography. The fractions obtained were analyzed by SDS-PAGE and Coomassie staining (right gel). The input (loaded material) and fractions 39 to 43 are shown. **B)** Count histogram of a mass photometry measurement of the FERRY complex. The peak for the fully assembled FERRY complex is indicated. **C)** Immunostaining of Fy-2 and Rabankyrin-5 in HeLa *wildtype* (*wt*) and *fy*-2 knock-out (KO) cells (upper panels) and Fy-4 and Rabankyrin-5 in HeLa *wt* and *fy*-4 KO cells (lower panels) (Scale bar: 10µm) (see also Methods: Antibody validation). **D)** Coomassie-stained SDS PAGE of an *in vitro* pulldown assay using GST, GST-Rab5:GDP and GST-Rab5:GTP to probe the interaction with the FERRY complex. **E)** Quantification of the co-localization of Fy-2 with Fy-4 and the endosomal markers EEA1 and Rabankyrin-5 as shown in C. **F)** Western blot analysis of HeLa Kyoto *wt* and different FERRY subunit KO HeLa cell lines against Fy-1 to Fy-5 and GAPDH as loading control.

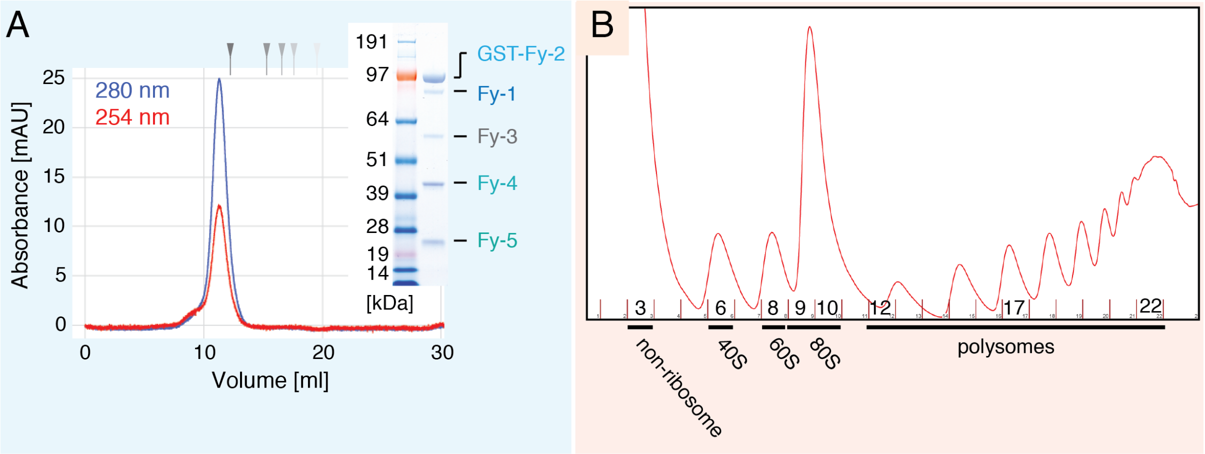


**Figure S2**: **A)** SEC profile of GST-FERRY (blue: 280 nm, red: 254 nm) with a Coomassie stained SDS PAGE of the peak fraction (inset). The grey arrows represent a molecular weight standard (670, 158, 44, 17, 1.35 kDa). **B)** Whole cell extracts prepared from HEK293 cell lines expressing 2x-Flag-PreScission-$His_6$ tagged Fy2 were separated by sucrose density gradient centrifugation (Absorbance profile (260 nm).

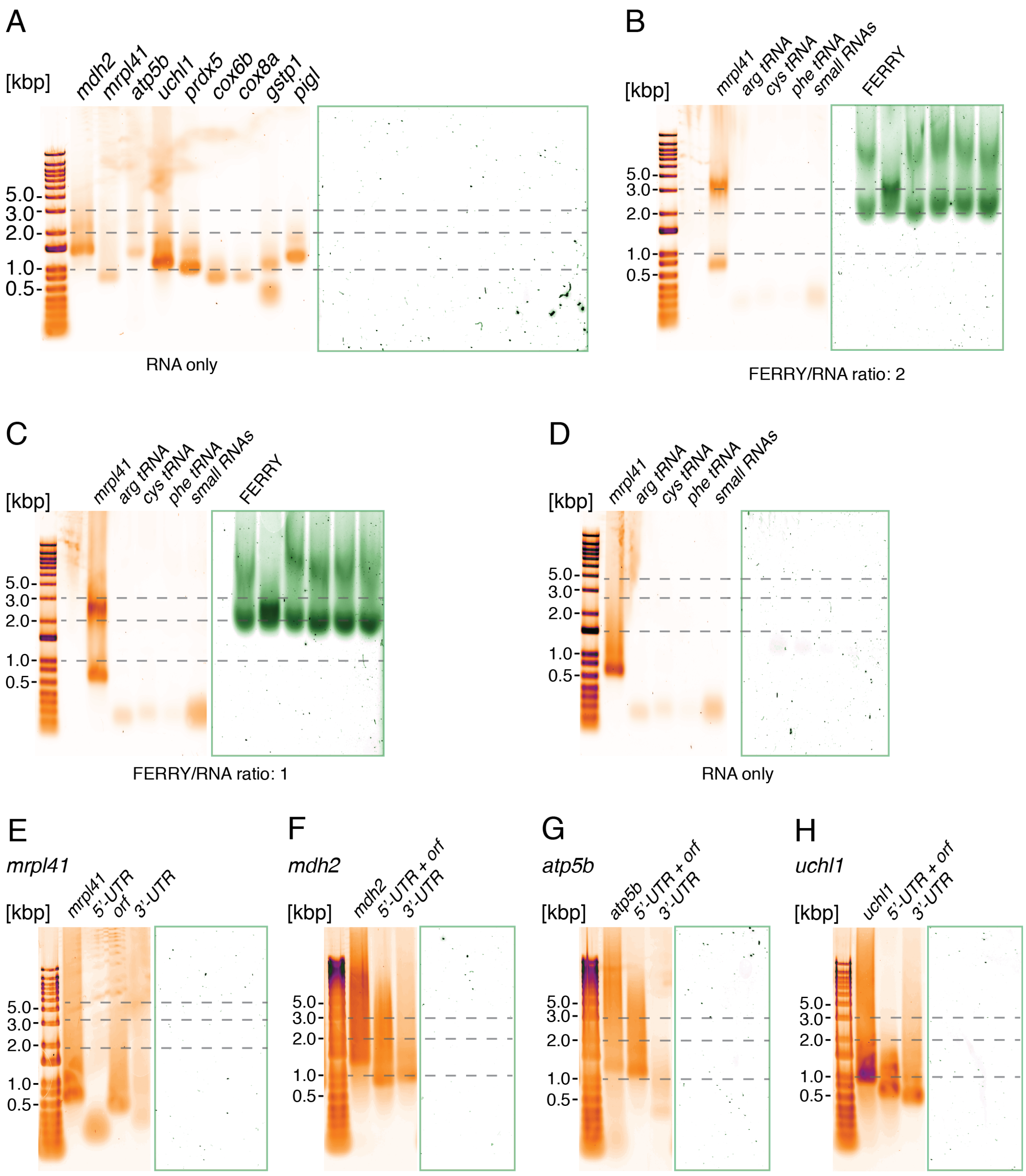


**Figure S3**: Electrophoretic mobility shift assay (EMSA) with different mRNAs in absence of the FERRY complex. (RNA: orange, SYBR Gold; proteins: green, Sypro Red). **B)** and **C)** EMSAs using *mrpl41*, three different tRNAs and an extract of small RNAs from HEK 293 cells at two different FERRY/RNA ratios. **D)** EMSAs using *mrpl41*, three different tRNAs and an extract of small RNAs from HEK 293 cells in the absence of FERRY. **E)** – **H)** EMSAs in the absence of the FERRY complex comparing four different RNAs with their respective subdivision construct shown in Figure 3E.

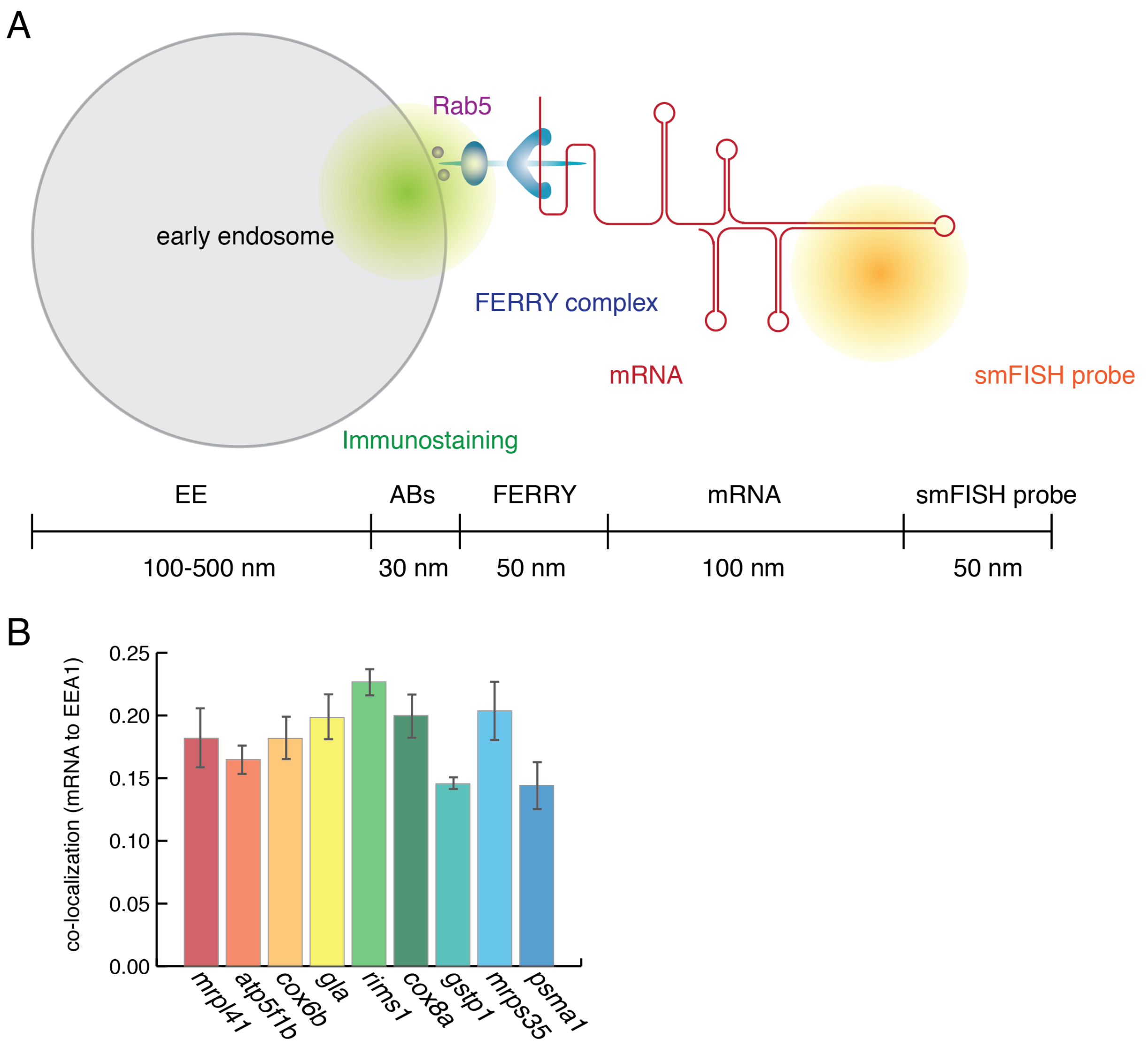


**Figure S4**: **A)** Schematic visualization and estimation of the scale of the early endosome (EE, grey), the FERRY complex (blue), EEA1 (green) and mRNA (orange) with the respective fluorescent labels either antibodies or smFISH. **B)** Quantification of the co-localization of different mRNAs to EEA1 in HeLa *wt* cells.

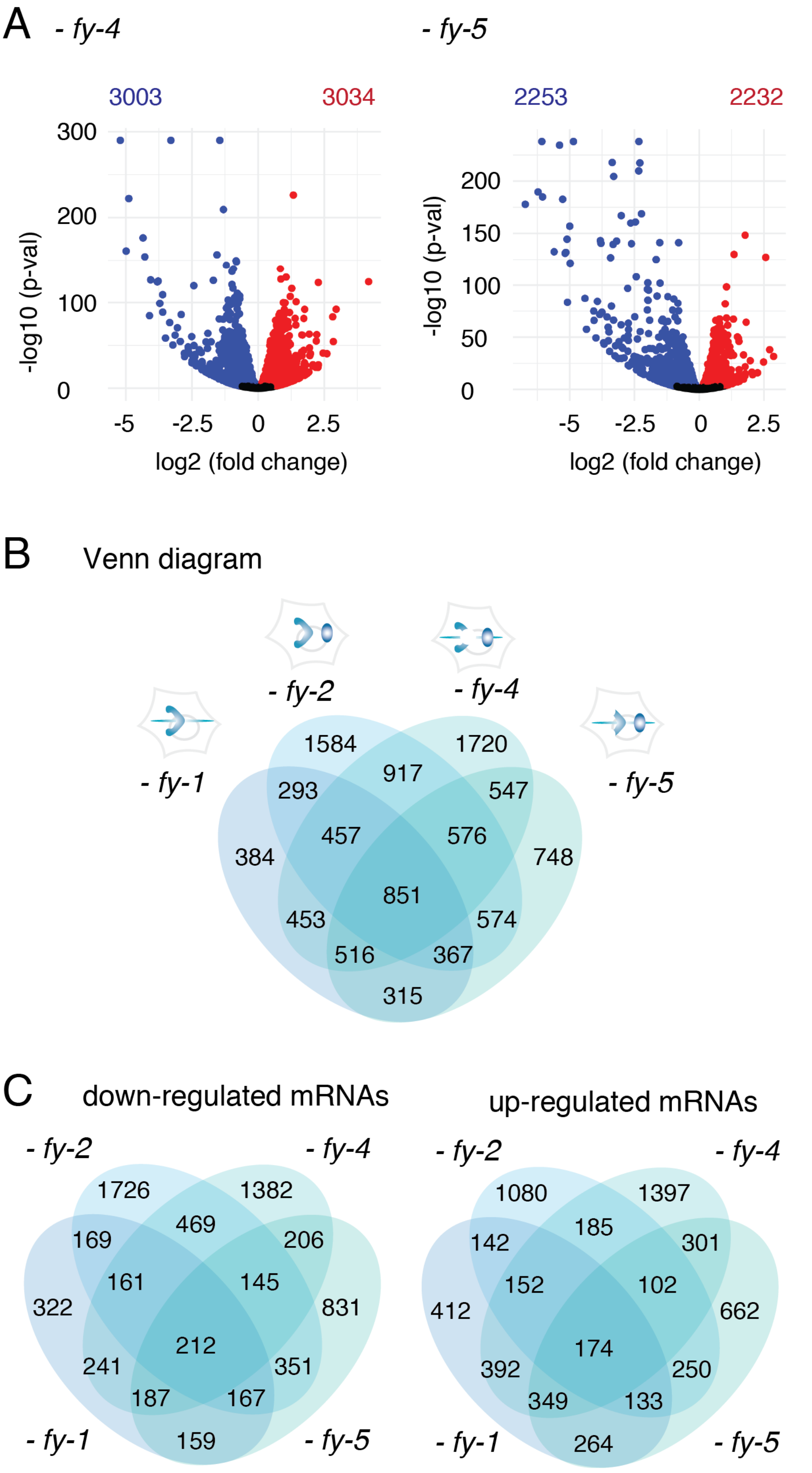


**Figure S5: A)** Volcano plots of the transcriptomic changes in the *fy-4* and *fy-5* KO cell lines compared to *wt*. (blue: down; red: up). **B)** Venn diagram of the transcriptome data of the four different FERRY component KO cell lines **C)** Venn diagrams of the FERRY KO cell lines only showing down-regulated (left) and up-regulated (right) genes.

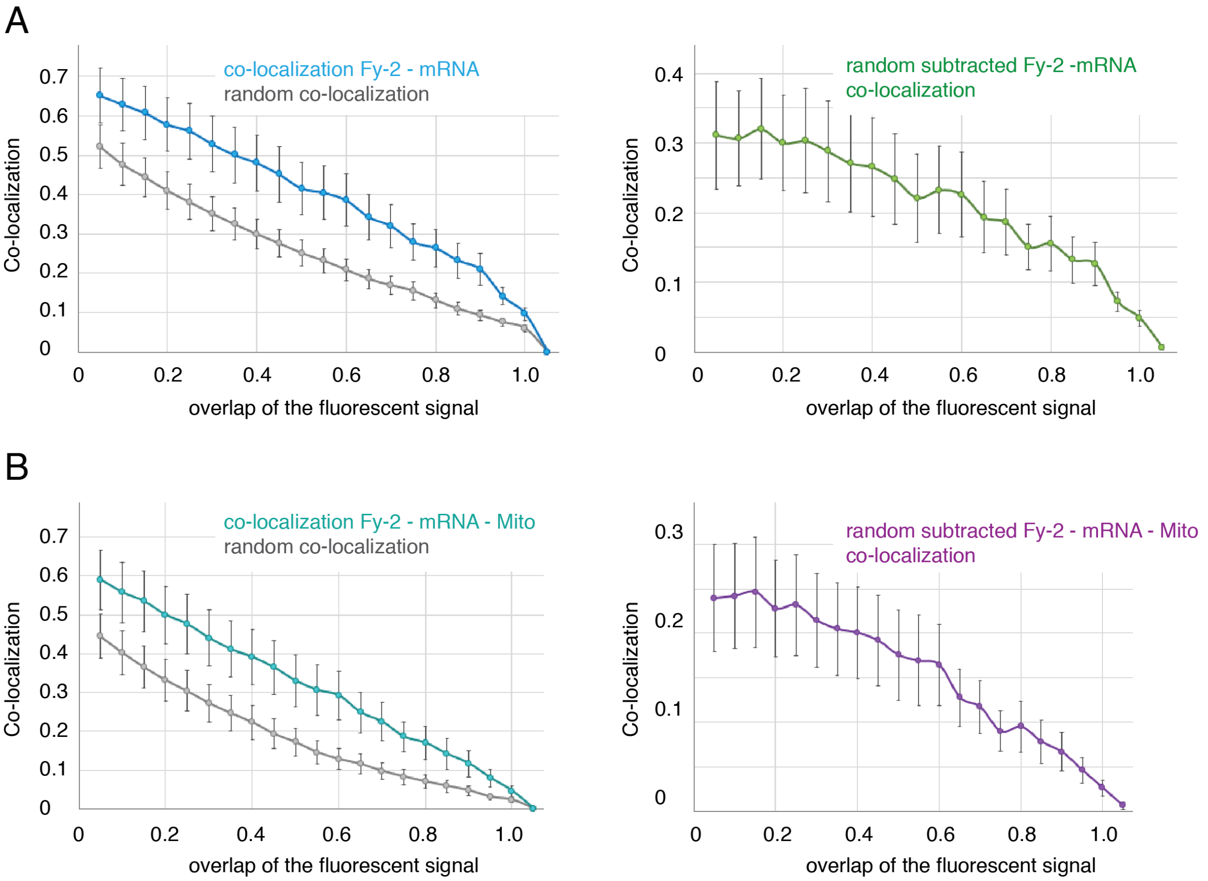


**Figure S6: A)** Left: Observed co-localization of *polyA* and Fy-2 (blue) compared to the calculated random co-localization (grey). Right: Subtraction of the random co-localization from the observed *polyA* - Fy-2 co-localization (green). **B)** Left: Observed co-localization of *polyA*, Fy-2 and mitochondria (turquoise) compared to the calculated random co-localization (grey). Right: Subtraction of the random co-localization from the observed *polyA* - Fy-2 - mitochondria co-localization (purple).
